# Network dynamics for sensory prioritization: Functional connectivity related to individual sensory weighting of vision versus proprioception during upper limb control

**DOI:** 10.1101/2025.07.31.667932

**Authors:** K. Folco, A. Pilacinski, H. Cheng, S. D. Newman, M. Wali, R. Babu, H. J. Block

**Affiliations:** Indiana University, Bloomington IN; Ruhr University Bochum, Bochum, Germany; University of Alabama, Tuscaloosa AL

## Abstract

Precise control of the hand requires the dynamic integration of visual and proprioceptive (body position sense) sensory cues with internal models, task goals, and motor plans. Individual differences in how visual and proprioceptive cues are weighted have been related to neural substrates, such as posterior parietal regions, yet the underlying neural dynamics are unclear. This study investigated the relationship between visuo-proprioceptive perception during a bimanual pointing task and the whole-brain network dynamics using resting-state functional magnetic resonance imaging (rsfMRI) with both atlas-based and individually-localized ROI seeds. Our results confirm the existence of individual sensory biases, and find they have systematic influences on functional connectivity. Activity between sensorimotor regions and default mode network (DMN) nodes was related to sensory weighting (e.g. individual degree of reliance on vision versus proprioception) and may reflect updating of internal models. The ventral premotor cortex (PMv) emerged as an important node with functional connections suggesting its role for integrating motor plans, internal models, and sensory percepts. Connections from multisensory integration regions like the middle temporal gyrus (MTG) and the superior parietal lobule (SPL), and motor coordination regions in the cerebellum, were related to increased reliance on visual versus proprioceptive information. These findings suggest that individual sensory biases during sensorimotor behavior may be characterized by specific and specialized patterns of neural activity. This research establishes a foundational exploration of the neural systems underlying sensorimotor processing, supporting theories suggesting that learning in either the sensory or motor system may also cause plasticity in the other.

## Introduction

The neural basis for individual differences in sensorimotor function has not been well established. Measures of functional connectivity have provided some insight into the intrinsic neural patterns unique to individuals, yet it is still not well understood how these network dynamics relate to behavior, or each other. In the last two decades, there has been a growing adoption in neural investigations of the theory of neural networks in the brain, which posits that specific patterns of neural activity involving diverse and fluid neural regions/systems are related to specific behavioral functions or states (Bassett & Sporns, 2017; Lungarella & Sporns, 2006; Sporns et al., 2004). Under this perspective, it may be necessary to consider the whole neural system when interrogating a specific behavioral output, such as estimating the hand’s location using sensory cues. Perceiving and localizing one’s hand is a critical component of upper limb movement and skilled behavior. This process is multisensory, involving both proprioceptive cues from the joints and muscles and visual cues from the eyes.

Sensory and motor functions are heavily integrated in human behavior and likely make use of similar interacting and diverse networks to process and engage with the environment (Boyd et al., 2024; S. Huang et al., 2015; Lungarella & Sporns, 2006; Vahdat et al., 2011a). Indeed, investigations of repeated practice with specific visual cues and motor movements (i.e. expertise) have related these experiences to altered sensory and motor processing, which suggested that individual sensory and motor experiences may alter sensory and motor processing (Charness et al., 2005; Fitch & Barnstaple, 2024; Hänggi et al., 2010; Reingold & Charness, 2005). There is also a growing body of research that has related early sensory and motor area activity to cross-system cues, such as primary motor (M1) neurons responding to somatosensory stimuli, or proprioceptive cues causing a distorted visual perception of angle (Bhalla & Proffitt, 1999; Carel et al., 2000; Ebrahimi & Ostry, 2024; Ghazanfar & Schroeder, 2006; Kumar et al., 2022; Lewis & Byblow, 2004; Mirdamadi et al., 2022a; Shaikhouni et al., 2013; Yau et al., 2015). These examples demonstrate that sensory and motor functions are heavily integrated, related directly to individual experience, and likely make use of a similar interacting distributed networks to process and engage with the environment (Boyd et al., 2024; S. Huang et al., 2015; Lungarella & Sporns, 2006; Vahdat et al., 2011a).

Many neural systems and regions have been identified as involved in estimating hand position. Some of the traditionally established regions related to motor control or visuo-motor recalibration are ventral premotor cortex (PMv) (Vahdat et al., 2011b, 2014), dorsal premotor cortex (PMd) (Genon et al., 2016; Pilacinski & Lindner, 2019), primary motor cortex (M1) (Mirdamadi et al., 2022b; Munoz-Rubke et al., 2017), primary somatosensory cortex (S1) (Ebrahimi & Ostry, 2024; Kumar et al., 2019; Nasir et al., 2013; Ostry et al., 2010), posterior parietal cortex (PPC) including superior parietal lobule (SPL) (Limanowski & Blankenburg, 2016a) and further divided by the intraparietal sulcus (IPS) (Grefkes & Fink, 2005a; Pilacinski & Lindner, 2019; Randerath et al., 2017), middle temporal gyrus (MTG, including the dorsal posterior portion MT+) (Antal et al., 2004; Gao et al., 2020), the lateral occipital cortex (LO) (Q. Huang et al., 2022; Lesser et al., 1998; Mullin & Steeves, 2011), and the cerebellum (Donchin et al., 2012; Galea et al., 2011). To complicate this extensive list, some of these regions have been identified as members of more than one neural network, indicating their potential as network hubs, or regions that communicate information between networks (Cole et al., 2013; Van Den Heuvel & Sporns, 2013). For a summary of relevant networks and regions related to hand perception, see Table 1. This complex interplay of multiple neural systems should be considered when developing models of motor control or perception.

**Table 1:**
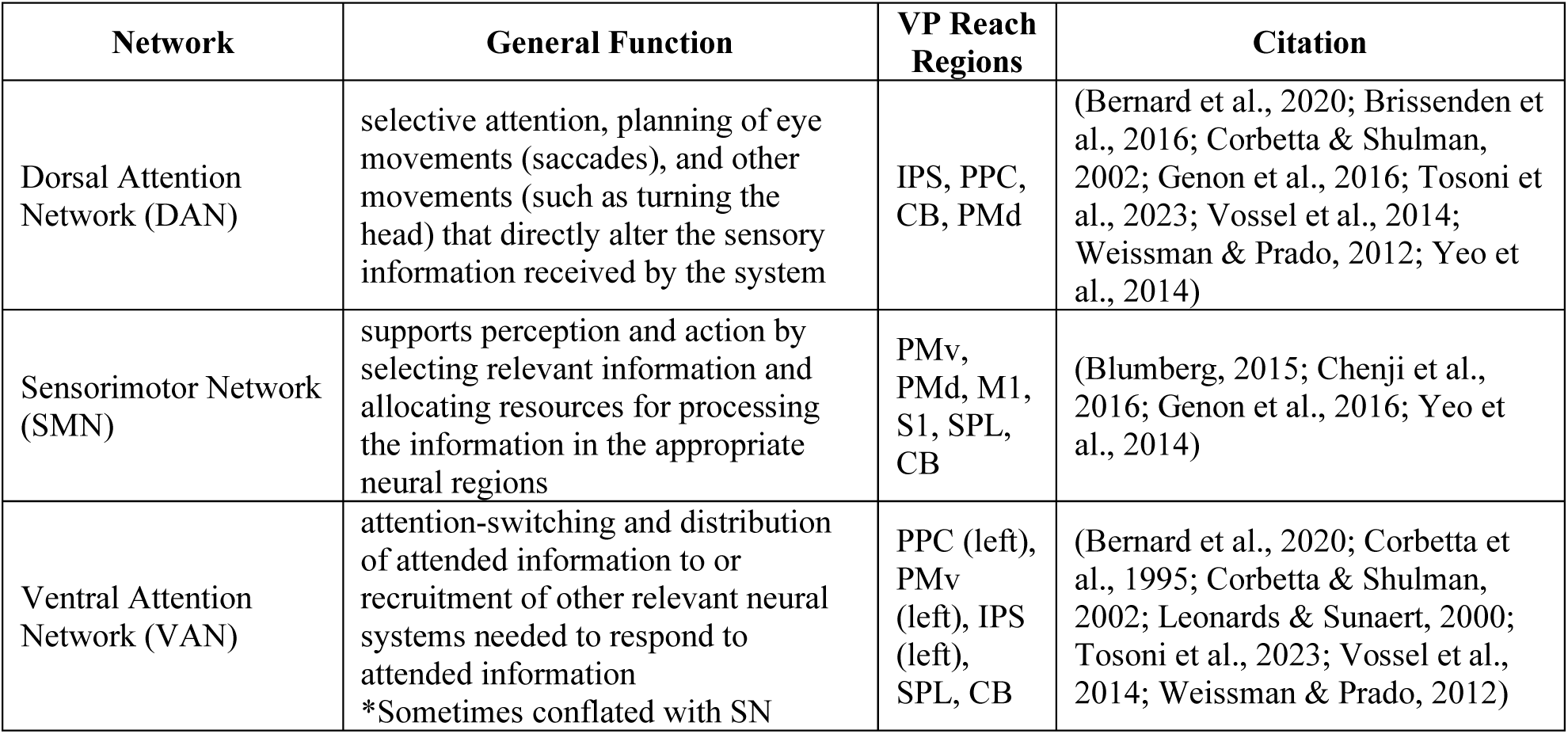
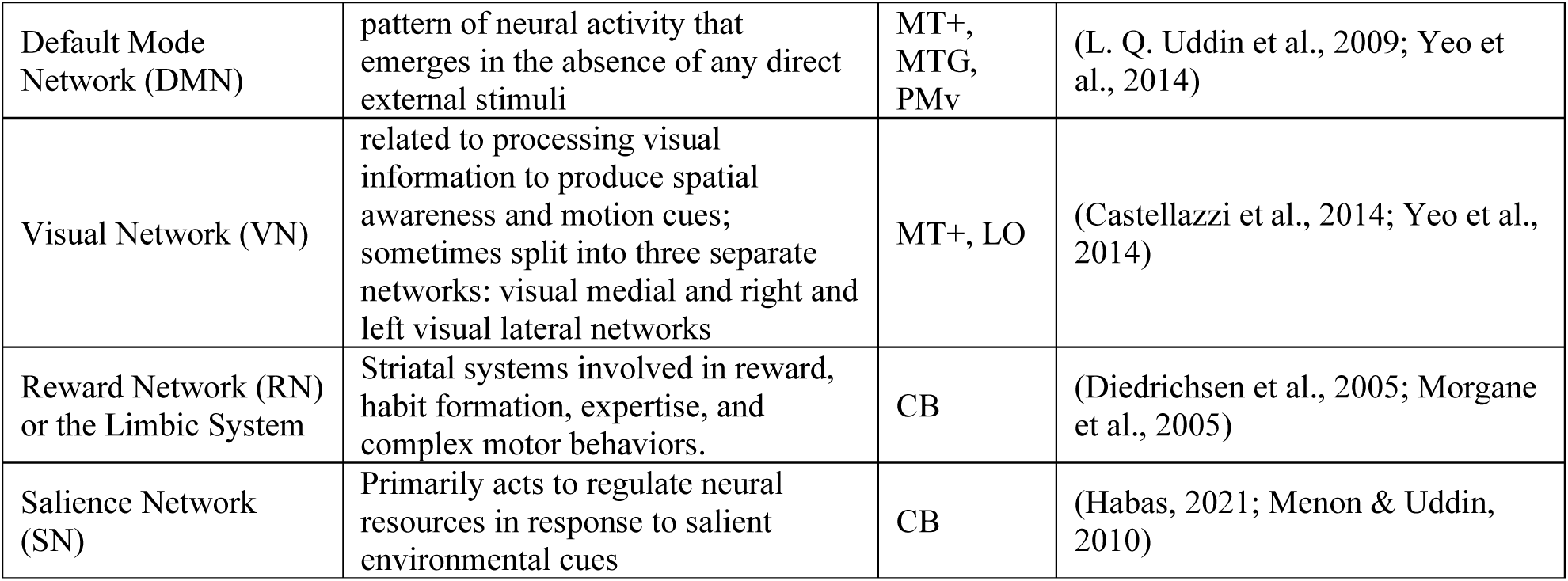
Neural networks associated with the ROIs related to visual-proprioceptive upper-limb reaches

Perhaps the most convincing case that hand perception involves the systematic coordination and co-evolution of multiple, seemingly disparate, brain networks, is the cerebellum. The cerebellum’s role in regulating network dynamics has been receiving increased attention, with an emerging status as a network hub region, or a region that is responsible for the integration, translation, and regulation of information between brain regions or networks (Kawabata et al., 2022). Functional connectivity between cerebellum lobules VI to VIII and the dorsal attention network (DAN) was related to cognitive task performance; cerebellum lobule VI (CB6) was related to visuospatial processing and attention; and cerebellar, visual, and striatal connectivity was related to movement disorder progression (Brissenden et al., 2016; H. Chen et al., 2022; Sako et al., 2021).

There is also a relationship between the cerebellum and drug reward, which is particularly interesting when considered alongside the shift during substance use from goal-driven behavior to more automatic or habitual striatal processing related to obtaining and using the addictive substance. These cerebro-cerebellar connections underscore the importance of applying a whole brain dynamic network approach and suggest the possibility of capturing individual differences in neural activity patterns.

Whole-brain functional connectivity can simultaneously capture widespread neural activity and regions that have been related to hand movements, which usually involve motor, visual, proprioceptive and multisensory systems. Individual neural activity patterns may be related to properties of an individual’s visual, proprioceptive, or multisensory processing. For example, a person may rely more heavily on vision than proprioception, and the degree to which vision versus proprioception is used when pointing to the estimated location of the other hand can be computed as a sensory weighting (Block & Bastian, 2011a).

We first hypothesized that behavioral measures of sensory behavior would be related to the neural system activity patterns measured as whole brain resting state functional connectivity. Specifically, we tested the relationship between resting state functional connectivity and four behavioral measures of sensory behavior. It was expected that two measures of sensory weighting, and one measure each of visual and proprioceptive precision (measured as amount of deviation of reach endpoint on each trial type) would be related to neural activity. Alternatively, neural connectivity may depend on motor performance overall, such as speed of reach or overall accuracy. Our second hypothesis was that there would be widespread recruitment of neural regions and systems previously identified as involved in multisensory hand perception. Specifically, this hypothesis makes the tentative assertion that early sensory and motor area activity, in addition to regions traditionally considered exclusively motor, will be related to sensory behavior, in line with a distributed systems model of sensorimotor processes. Using six neural regions as initial guidance, we tested either the regions themselves and/or the networks they had previously been associated with, under the expectation that this would provide a comprehensive picture of neural dynamics during hand localization. Alternatively, it may be the case that this function relies on more focal and localized neural activity.

More specifically, we expected key hub regions such as the cerebellum to have multiple connections related to individual sensory behavior, with functional correlations between CB and regions in the sensorimotor network (SMN), attention networks (including the salience network, SN), visual networks, and striatal networks. In addition, multisensory integration regions such as SPL, and sensorimotor regions such as PMv, should have functional connections to attention network(s) and the SMN. We also expect unisensory regions such as S1 and LO to demonstrate functional connectivity differences related to individual sensory behavior with regions in the SMN, attention networks, and visual networks. Finally, we expected that motor regions such as M1, which has previously been identified as responsive to sensory stimulation (Mirdamadi et al., 2022b; Munoz-Rubke et al., 2017), would show functional connections with sensorimotor or sensory regions that were related to sensory behavior.

## Methods

### Participants

Initial screening resulted in 70 participants. Six individuals were excluded for failure to perform the reaching task correctly, as measured by tendency to reach toward fixation rather than target. One was excluded due to suspected intoxication during the study which caused extreme difficulty in following directions. One person stopped responding after the first session, and their data was excluded. An additional 8 participants were excluded due to technical errors that resulted in incomplete data collection. This left 55 participants that were used for subsequent analysis. These participants were right-handed adults, 39 female, mean age of 23.93 ± 1.79, located in the Bloomington Indiana area. All provided written informed consent under the Institutional Review Board (IRB) protocol number 13138 and received payment for their participation in the form of Kroger or Amazon gift cards. The racial and ethnicity distributions of our participants included 45.45% White non-Hispanic or Latino, 7.27% White Hispanic or Latino, 32.72% Asian, 3.64% Black or African-American, 5.45% were more than one race and not Hispanic or Latino, and 5.45% were more than one race and Hispanic or Latino.

All participants reported having normal or corrected-to-normal vision; being free of neurological or orthopedic or pain conditions, claustrophobia, or difficulty remaining still; and being free of metallic, mechanical, or magnetic implants. Women who were pregnant or thought they might have been pregnant were also excluded, as were women using an intrauterine device that might affect MR compatibility.

### Familiarization procedure

Participants completed a familiarization and a main session on different days. During familiarization, participants completed consent and screening forms and read more detailed information about the procedures and tasks in a self-paced PowerPoint presentation. They next completed 8 practice trials of the pointing task, followed by a block of 40 trials to ensure they could follow task instructions (see “Pointing task” below).

Eligible participants were then introduced to the procedure for magnetic resonance imaging (MRI) using the mock MRI located in the imaging facility. Participants laid supine, head first, in the mock MRI and were slid into the bore. The lights were then turned off and standard MRI sounds were played on a speaker system in the mock MRI for about 10 minutes to best mimic the environment of the MRI. Participants then received instructions on how to perform the functional localizer task used in the main session (see “Functional localizer” below). This concluded the familiarization session. Participants who could tolerate the MRI environment and follow task instructions were eligible for the main session.

During the main session, participants again practiced 8 trials of the pointing task to refresh their memory. Next, they completed the MRI safety screening form and the MRI technician confirmed eligibility. Participants were then escorted to the MRI, given hearing protection, and laid supine in the bore head first. In the first MRI scan session, diffusion images (∼3 minutes) and 12 minutes of resting state functional imaging (rsfMRI) data was collected, referred to as the “Pre” rsfMRI condition. Following that scan sequence, participants relocated to the behavioral testing room down the hall and performed 73 trials of the pointing task which took about 15 minutes. This was followed directly by another 12 minute rsfMRI sequence, “Post” rsfMRI condition. As part of a separate study, 36 participants were again removed from the MRI and completed an additional block of the pointing task (73 trials) and another 12 minute rsfMRI sequence. Directly following the final rsfMRI sequence, two blocks of functional imaging were completed to localize the right M1 and S1 (see “Functional localizer” below). Two runs (∼4 minutes each) of functional MRI were collected for the localizer analysis. After this final fMRI scan sequence, the experimenter exited the scan room and participants were instructed lay still for a few more minutes for their anatomical scan (∼7 minutes) and another diffusion sequence (∼3 minutes). The Pre/Post resting state data was the focus of the current report, and the localizer was used to identify subject-specific regions.

Upon completion of the last MRI scan sequence, participants completed an electronic debriefing questionnaire that assessed general mood and thought processes for separate analysis not reported here. Participants were then given a copy of their anatomical scan, debriefed, and paid for their participation.

### Pointing Task

#### Apparatus

Outside the scanning room, a 2D virtual reality apparatus was used which had two-sided touchscreen (PQLabs), mirror, and LCD screen. These were all positioned in the horizontal plane. Participants viewed the task display in the mirror, which reflected the LCD screen. These images appeared to the participant to be in the plane of the touchscreens, below the mirror (Fig 1). No direct vision of the hands was possible, and black fabric was used to obscure the participant’s view of their arms and shoulders.

**Figure 1:**
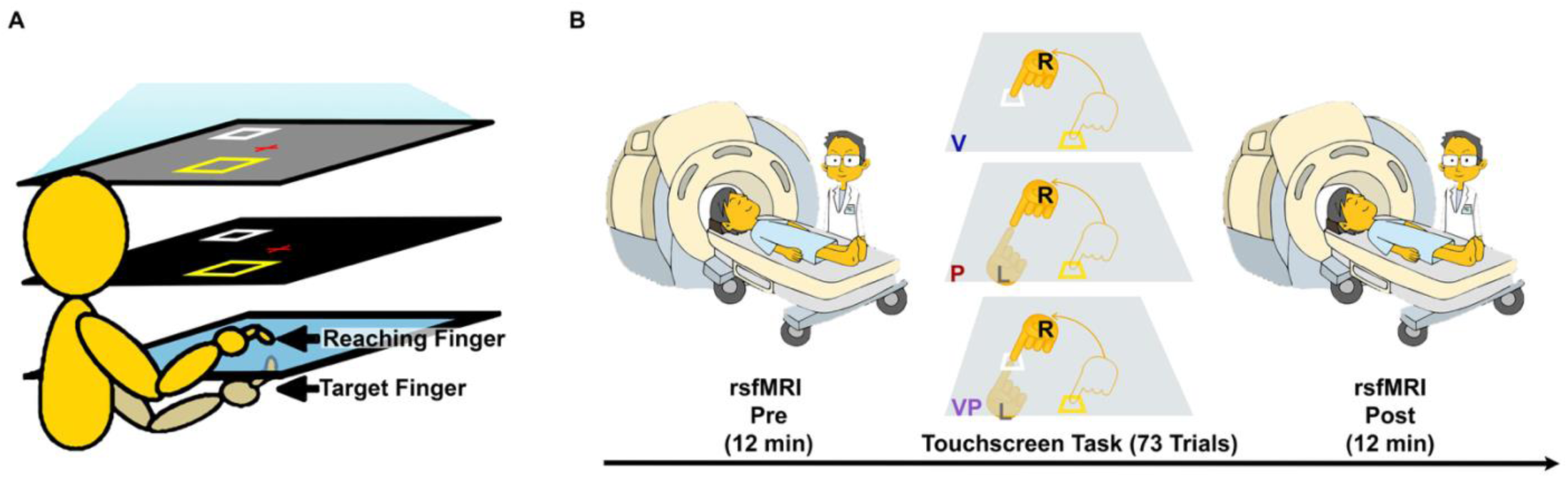
Pointing task and procedure. Panel A: Apparatus showing the right hand as the reaching to the white box V target. Top square is monitor, middle square is mirrored display that participant sees, and blue box is the double-sided touch screen. Panel B: Experimental procedure for the main session showing the two MRI scan sessions with the reaching task in between. The three trial types, V (visual), P (proprioceptive), VP (visual-proprioceptive) are shown for the touchscreen task, R indicates right hand, L left, arrows for reach trajectory.

**Figure 2:**
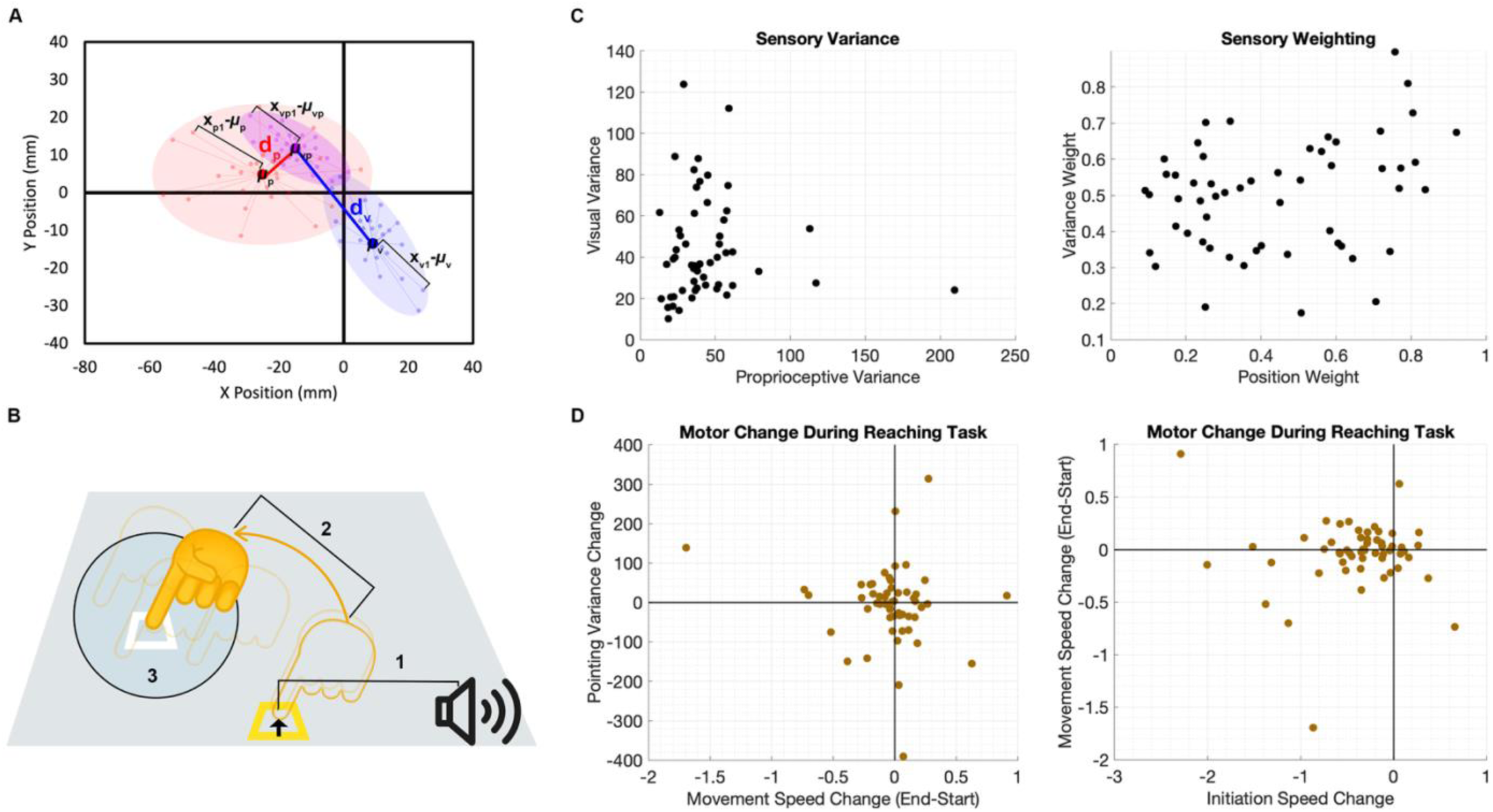
Visualization of the measurement and distribution of individual behavioral measures in our sample. Panel A: Sensory computations, where x refers to individual reach endpoints, µ refers to the mean reach endpoint for a given reach condition, d refers to the distance between the mean endpoint for each condition, and subscripts refer to the type of reach target (v: visual, blue, p: proprioceptive, red, vp: visual and proprioceptive, purple). Panel B: Motor computations, where 1 is initiation speed, 2 is movement speed, 3 is pointing variance. Panel C: Subject distribution of sensory measures, note the unique distributions for all four measures. Panel D: Subject distribution of motor measures, note there were few subjects who improved speed or precision (points are clustered around zero).

**Figure 3:**
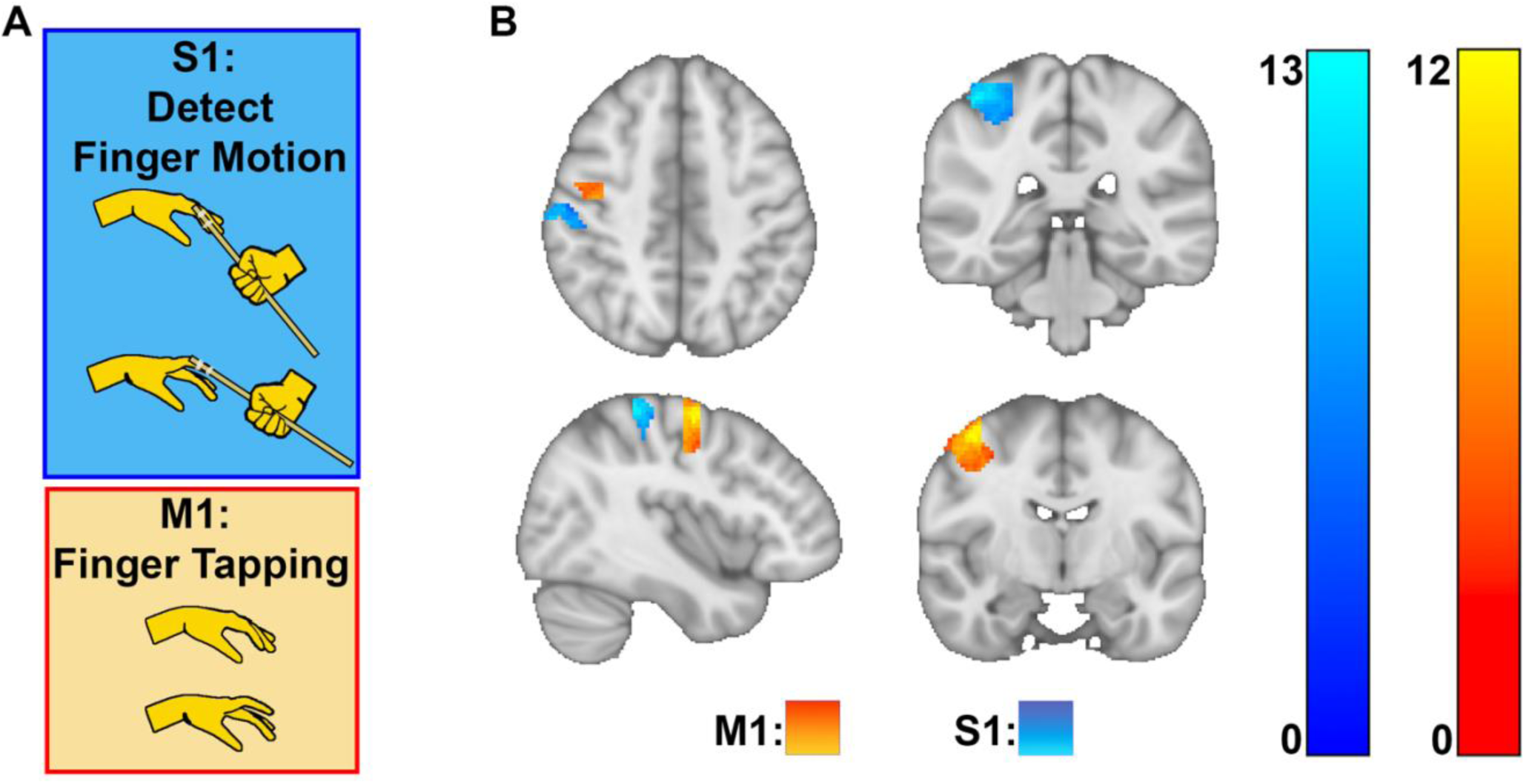
Panel A: Localizer tasks for S1 (blue) and M1 (orange). Panel B: Heatmap of averaged resulting ROIs with light blue or yellow indicating the most overlap across participants, and dark blue or red indicating the least participants.

#### Stimuli

Visual and proprioceptive cues acted as reach targets. Participants were asked to use their right “indicator” finger to reach and point on the upper touchscreen to where they perceived the reach target to be located. There were three target types: visual cue only (V), proprioceptive cue only (P), and multisensory (visual and proprioceptive, VP). The proprioceptive cue for P and VP trials was the participant’s left index fingertip positioned on a tactile marker on the lower touchscreen. The visual cue for V and VP trials was a 12 mm white square. During V trials, the left hand rested in the participant’s lap. During VP trials, the visual cue was displayed veridically over the proprioceptive cue.

#### Single Trial Procedure

At the start of each trial, a yellow square appeared in one of five possible positions arranged in a plus sign at the participant’s midline. To begin the trial, participants placed their indicator finger in the yellow square on the upper touchscreen. To help participants achieve the starting position, visual feedback as a blue circle (8mm) of the indicator finger near the start box was provided. It disappeared once the reaching finger was confirmed inside the start box. A red fixation cross appeared at random coordinates within 10 cm of the target and the participants were instructed to keep their eyes on the cross. Finally, a “go” signal (an auditory chime) instructed them to begin the trial, followed by pre-recorded spoken directions. They were asked to indicate the target position with as much accuracy as possible and were given no speed restrictions. The participants were asked to lift their indicator finger off the glass and place it down only at the perceived position of the target.

They were instructed not the drag their finger on the touchscreen. The participants had to hold on to the perceived position for 2s for the system to register their endpoint position, interpreted as their estimate of the target’s location. The trial was indicated to be complete by a camera-click sound, at which time the participant was instructed to remove all contact with the touchscreen so the next trial could begin. The participants received no performance feedback or knowledge of results, making sure that they had no way of knowing the accuracy of their performance.

Targets were presented at one of two possible locations roughly 10 cm ahead of the start positions and 4 cm apart from each other. To make sure that the participants could not memorize movement extent or direction, the start and target positions were randomized across trials. During the main session, participants performed a block of 73 pointing trials in between the “pre” and “post” rsfMRI scans: 37 VP, 18 P, and 18 V trials. These were presented in a repeating order (VP-P-VP-V).

### Analysis of pointing data

#### Unimodal variance

We estimated the two-dimensional variance associated with each unimodal estimate, visual and proprioceptive, from the indicator finger’s endpoint position on the unimodal V and P trials, respectively. These estimates are assumed to include motor and proprioceptive noise from the indicator finger rather than pure sensory variance. For each set of endpoints, a vector was computed from the mean endpoint position to each individual endpoint. The variance of the vector magnitudes was computed as an approximation of visual or proprioceptive estimation variance (σ_v_ ^2^ and σ ^2^).

#### Variance weighting

To estimate how much participants should be relying on vision vs. proprioception when pointing to VP cues if they are minimizing variance, we computed a ratio of the unimodal variances (Formula 1). Variance weight can range from 0 (only relying on proprioception) to 1 (only relying on vision), with 0.5 suggesting equal reliance on vision and proprioception due to equivalent variances. In other words, if σ ^2^ is much smaller than σ ^2^, multisensory integration should favor proprioception, and the variance weight should be less than 0.5.

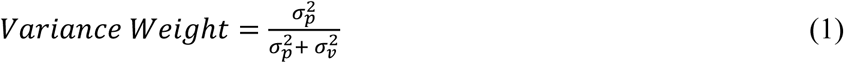

#### Position weighting

We also computed an estimate of visuo-proprioceptive weighting based on positional biases rather than variance (Formula 2). This method takes advantage of participants’ naturally different spatial biases when estimating visual vs. proprioceptive cues with no feedback (Crowe et al., 1987; Foley & Held, 1972). For each VP trial, we computed the 2D distance from the indicator finger endpoint to the mean of the 4 closest V trials in the sequence (d_v_ – d_vp_) and the mean of the 4 closest P trials in the sequence (d_p_ – d_vp_). We computed a ratio of these distances for each VP trial, averaging the result across trials to obtain position weight:

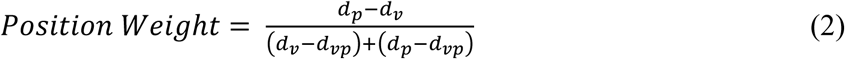

Position weight can range from 0 (only relying on proprioception) to 1 (only relying on vision), with 0.5 suggesting equal reliance. Unlike variance weight, position weight depends on where the VP estimate falls in relation to the V and P estimates. In other words, if a person’s VP estimate is closer to where they estimate the P cue than to where they estimate the V cue on unimodal trials, we interpret this as the person relying more on proprioception than vision when both are available (position weight < 0.5). Unlike variance weight, position weight reflects the participant’s actual performance when estimating VP targets and may be influenced by factors other than variance, such as task goals and attention (Block & Bastian, 2011b; Block & Sexton, 2020).

#### Motor control measures

Three measures of motor behavior were also computed to confirm there was no motor learning from performing reaches without feedback. It was also of interest whether there were any differences in individual motor abilities. First, initiation speed was computed as the duration between the auditory reach cue (signaling that the participant should begin their reach) and the moment the participant lifted their indicator finger from the touchscreen to begin their reach. Second, movement time was computed as the duration of the reaching movement, that is, how long in between lifting their indicator finger off the touchscreen from the start position and placing it back onto the touchscreen at the estimated target location. Lastly, pointing variance was the computed variance of indicator finger endpoints across trial types. To reflect whether these variables underwent any change from early to late in the task, we subtracted the first four trials from the last four trials, resulting in a single value per participant.

### Functional Localizer Task Procedure

An experimenter entered the MRI scan room and the participant was slid out of the bore, but they remained supine on the bed. They were verbally reminded of the directions while the experimenter used body tape to secure a 3D printed rod to the left index finger of the participant. The rod was attached at one end to a 36 inch wooden dowel that extended outside the bore. The participant was given a button-box and it was confirmed their right index finger and right middle finger were aligned with the correct buttons. The participant was then re-entered into the MRI bore. The experimenter donned headphones and remained inside the scan room with the participant. A second experimenter controlled a Matlab program from the control room using psychtoolbox to present audio cues to the experimenter inside the MRI room. There were three task blocks repeated six times each in total, presented in a random order. During the rest block, nothing occurred, and this was used as a baseline condition.

The two localizer tasks were designed to target the M1 and the S1. To identify the subject’s M1, the participant was instructed to move their left index finger each time they felt the experimenter tap their left shin. The experimenter tapped their left shin every 2 seconds, responding to an audio cue heard over the headphones from the task program. The participant alternated moving their finger first up, then down. To identify the participant’s S1, the experimenter moved the participant’s left index finger either up or down using the dowel apparatus depending on the audio cue the experimenter heard (randomized, but evenly occurring). To ensure the participant was paying attention to how their finger was being moved, they were instructed to press their right index finger on the button when their left index finger was moved up, and their right middle finger when their left index finger was moved down. This data was recorded but not analyzed beyond confirming that participants had followed task instructions. The 18 total blocks (6 rest, 6 M1, 6 S1), were broken into two functional runs that each lasted 4 minutes. Timing signals from the audio cues sent to the experimenter were synced with the MRI pulse sequence and were used for subsequent analyses.

### MRI data acquisition

All MRI images were collected on a Siemens 3 Tesla whole body MRI scanner (Magnetom Prisma fit), with participants laying supine in the bore headfirst, using a 64 channel head coil, located in the Imaging Research Facility in the Psychology Building at Indiana University. The scan protocol began with diffusion images (∼3 minutes), followed by a 12 minute resting state sequence. After an intermission for the behavioral task, another 12 minute resting state sequence was collected. Following a final behavioral task intermission, the final resting state sequence was collected (12 minutes), in addition to the anatomical scan sequence (5 minutes). A functional localizer task was also performed (∼8 minutes) for the regions of S1 and M1. As part of a separate study question, a second set of diffusion images were also collected (∼3 minutes).

### Functional localizer imaging parameters

The functional MRI event-related images were acquired across two scans, each having a total of 240 volumes collected in a tranverse plane using a T2-weighted gradient-echo echo-planar-imaging (EPI) sequence with the following parameters: repetition time (TR) of 1.14 s, echo time (TE) of .03, flip angle of 60 °, field of view (FOV) of 216 x 216 mm, matrix size of 108 x 108, slice thickness of 2 mm, spacing between slices of 2 mm, 72 slices, and voxel size of 2 x 2 x 2 mm^3^.

### Resting state functional imaging parameters

The functional MRI images collected had a total of 1000 volumes collected in a transverse plane using a gradient-echo echo-planar-imaging (EPI) sequence with simultaneous multi-slice (SMS) acquisition with the following parameters: repetition time (TR) of .72 s, echo time (TE) of .03 s, flip angle of 52°, field of view (FOV) of 216 x 216 mm, matrix size of 108 x 108, slice thickness of 2 mm, spacing between slices of 2 mm, 72 slices, and voxel size of 2 x 2 x 2 mm^3^. The SMS factor was 8.

### Diffusion tensor imaging parameters

The DTI sequence included 92 slices acquired using diffusion weighted spin-echo single-shot planar imaging applying a pair of diffusion gradients in the anterior-posterior direction with b-values of 1000 s/mm^2^ implemented in 30 directions, including four volumes with no diffusion gradient, b = 0 s/mm^2^. Other parameters are as follows: FOV of 222 x 222 mm, matrix size of 148 x 148, slice thickness of 1.5 mm, spacing between slices of 1.5 mm, an echo time (TE) of .08 s, and a repetition time (TR) of 3.4 s, resulting in a voxel resolution of 1.5 x 1.5 x 1.5 mm^3^.

## MRI Analysis

### Distortion correction

FSL software was used to create and apply field maps to functional images (Jenkinson et al., 2012). Field maps were created from two DTI images with opposite phase-encoding directions that had b-values of 0 s/mm^2^ (b0 images) that were combined into a single volume using fslmerge. Specifically, FSL’s fslroi was used to extract a single volume from each diffusion image gradient, then FSL’s fslmerge was used to combine both gradients. Next, FSL’s topup’s --iout produced a field map which was then transformed with FSL flirt to the subjects’ functional image space using the mean functional image in native space. Next, the field map was applied to the functional images using FSL fugue with a dwell parameter of .00007250112, computed from the division of the effective echo spacing of .000580009 by the multiband acceleration factor of 8. The resulting distortion-corrected functional image was then used for subsequent analysis of the functional images. There was one subject whose diffusion images were not collected properly and therefore distortion correction was limited to the SPM12 realign & unwarp procedure (Andersson et al., 2001), which uses head movement to estimate derivatives of the deformation field then resamples functional data to align with the deformation field of the reference image.

### Functional Localizer

Event-related fMRI analyses were performed using the Statistical Parametric Mapping (SPM12) toolbox in Matlab 2023b (Ashburner et al., 2014; The MathWorks Inc., 2023). Also, the toolbox SPM_SS (Nieto-Castanon & Fedorenko, 2012) was used to perform the Group-constrained subject-specific (GcSS) analysis.

**Preprocessing**. Preprocessing was done using the SPM toolbox for Matlab with a pipeline involving realignment (to the first slice), slice timing correction, coregistration, segmentation, and normalization. Data were visually inspected for **artifacts** and the procedure was redone after resetting the origin if any errors were found. Functional data were realigned using SPM realign & unwarp procedure (Andersson et al., 2001), where all scans were coregistered to a reference image (first scan of the first session) using a least squares approach and a 6 parameter (rigid body) transformation (Friston et al., 1995), and resampled using b-spline interpolation to correct for motion and magnetic susceptibility interactions. Temporal misalignment between different slices of the functional data (acquired in interleaved order) was corrected with SPM slice-timing correction (STC) procedure (Henson et al., 1999; Whitfield-Gabrieli et al., 2011), using sinc temporal interpolation to resample each slice to a common mid-acquisition time. Functional and anatomical data were normalized into standard MNI space, segmented into grey matter, white matter, and CSF tissue classes, and resampled to 2 mm isotropic voxels following a direct normalization procedure (Calhoun et al., 2017; Nieto-Castanon, n.d.) using SPM unified segmentation and normalization algorithm (Ashburner, 2007; Ashburner & Friston, 2005) with the default IXI-549 tissue probability map template. Last, functional data were smoothed using spatial convolution with a Gaussian kernel of 8 mm full width half maximum (FWHM). Three task conditions were identified: rest, finger tapping, and finger sensing, and these times were convolved with the hemodynamic response function and estimated across all blocks (each condition had 6 total blocks).

**First-Level Analysis**. To perform the second-level analyses, we used the SPM_SS toolbox to compute the ROI masks from the overlapping activations across subjects. We tested the contrast of our functional condition against rest within each individual subject (e.g. finger tapping – rest = motor localizer M1, finger sensing – rest = proprioceptive localizer S1). There was large variability in functional activation clusters across subjects, emphasizing the importance of using this localizer task.

Second-Level Analysis. This section describes the analysis procedures used for the resting state functional data.

**Masking.** We used two masks to select relevant functional data from each participant to minimize the noise in the functional signal. First, we used the probabilistic Harvard-Oxford atlas ROIs of precentral and postcentral gyrus. We binarized, right-lateralized, and applied a 16% threshold to the probabilistic atlas ROI mask. To add more systematic specificity, we also used the results of an algorithmic procedure, GcSS, as a mask. Finally, only functional activity within both the atlas-mask and the GcSS mask was retained to define the subject-specific region.

**GcSS Analysis**. All 55 subjects informed the GcSS procedure. The GcSS procedure uses available data to generate regions of task-related functional activation that are common across the population, providing a more replicable and generalizable method for functional localization. The analysis was a subject-specific ROI-based (automatically defined by the package) analysis using a one-sample t-test with restricted maximum likelihood. We found it was necessary to establish four significance levels that permitted appropriate thresholds for each participant’s functional activation (Localizer contrast threshold of FDR-corrected p-value < 0.0001, FDR-corrected p-value < .05, uncorrected p-value < .0001, and uncorrected p-value < .05). All analyses used a threshold of 0.1 for the minimal voxel-level overlap, 0.5 for the minimal ROI-level overlap, and a smoothing factor of 8 mm. The most conservative significance level was the default (FDR-corrected p-value < 0.0001), and less conservative parameters were only explored when there were less than ten voxels of functional activation present in the masked region for a given subject. For the M1-localizer, we tested the contrast between M1-task blocks (finger tapping) and rest blocks to determine individual functional activation. For the S1-localizer, we tested the contrast between S1-task blocks (proprioception) and rest blocks to determine functional activation in each subject. As stated previously, an atlas mask was used to constrain the GcSS results, producing a single activation cluster within the Harvard-Oxford Atlas ROIs of either the right postcentral gyrus (S1) or the right precentral Gyrus (M1).

### rsfMRI

Results included in this manuscript come from analyses performed using CONN (Whitfield-Gabrieli & Nieto-Castanon, 2012) release 22.a (Nieto-Castanon & Whitfield-Gabrieli, 2022) and SPM (Penny et al., 2011)(RRID:SCR_007037) release 12.7771.

**Preprocessing.** Functional and anatomical data were preprocessed using a flexible preprocessing pipeline (Nieto-Castanon, 2020) including realignment with correction of susceptibility distortion interactions, slice timing correction, outlier detection, direct segmentation and MNI-space normalization, and smoothing.

Functional data were realigned using SPM realign & unwarp procedure (Andersson et al., 2001), where all scans were coregistered to a reference image (first scan of the first session) using a least squares approach and a 6 parameter (rigid body) transformation (Friston et al., 1995), and resampled using b-spline interpolation to correct for motion and magnetic susceptibility interactions. Temporal misalignment between different slices of the functional data (acquired in interleaved Siemens order) was corrected following SPM slice-timing correction (STC) procedure (Henson et al., 1999; Whitfield-Gabrieli et al., 2011), using sinc temporal interpolation to resample each slice BOLD timeseries to a common mid-acquisition time. Potential outlier scans were identified using ART (Whitfield-Gabrieli et al., 2011) as acquisitions with framewise displacement above 0.9 mm or global BOLD signal changes above 5 standard deviations (Nieto-Castanon, n.d.; Power et al., 2014), and a reference BOLD image was computed for each subject by averaging all scans excluding outliers. Functional and anatomical data were normalized into standard MNI space, segmented into grey matter, white matter, and CSF tissue classes, and resampled to 2 mm isotropic voxels following a direct normalization procedure (Calhoun et al., 2017; Nieto-Castanon, n.d.) using SPM unified segmentation and normalization algorithm (Ashburner, 2007; Ashburner & Friston, 2005) with the default IXI-549 tissue probability map template. Last, functional data were smoothed using spatial convolution with a Gaussian kernel of 8 mm full width half maximum (FWHM).

**Denoising.** In addition, functional data were denoised using a standard denoising pipeline (Nieto-Castanon, 2020) including the regression of potential confounding effects characterized by white matter timeseries (5 CompCor noise components), CSF timeseries (5 CompCor noise components), motion parameters and their first order derivatives (12 factors) (Friston et al., 1996), outlier scans (below 98 factors) (Power et al., 2014), session and task effects and their first order derivatives (6 factors), and linear trends (2 factors) within each functional run, followed by bandpass filtering of the BOLD timeseries (Hallquist et al., 2013) between 0.008 Hz and 0.09 Hz.

CompCor (Behzadi et al., 2007; Chai et al., 2012) noise components within white matter and CSF were estimated by computing the average BOLD signal as well as the largest principal components orthogonal to the BOLD average, motion parameters, and outlier scans within each subject’s eroded segmentation masks. From the number of noise terms included in this denoising strategy, the effective degrees of freedom of the BOLD signal after denoising were estimated to range from 322.5 to 343.6 (average 339.4) across all subjects (Nieto-Castanon, n.d.).

**First-Level Analysis**. Connectivity is computed at the individual subject level, estimated characterizing the patterns of functional connectivity with 164 HPC-ICA networks (Nieto-Castanon & Whitfield-Gabrieli, 2022), Harvard-Oxford atlas ROIs (Desikan et al., 2006), and the cerebellar parcellation from the Automated Anatomical Labeling (AAL) atlas (Tzourio-Mazoyer et al., 2002), and the two localizer ROIs for right M1 and S1. Functional connectivity strength was represented by Fisher-transformed bivariate correlation coefficients from a weighted general linear model, weighted-GLM (Nieto-Castanon, 2020), defined separately for each pair of seed and target areas, modeling the association between their BOLD signal timeseries. Individual scans were weighted by a boxcar signal characterizing each individual experimental condition (e.g. the resting state scans collected before and after reaching task making the “pre” and “post” conditions) convolved with an SPM canonical hemodynamic response function and rectified.

***Seed-Based Connectivity Maps***. Individual subject seed-based connectivity maps (SBC) contain the functional connectivity between every ROI parcel and every voxel in the brain, computed as the bivariate Fisher-transformed correlation coefficients between an ROI BOLD timeseries and the timeseries of each individual voxel. For subsequent analyses, we selected a total of 25 seeds, chosen for specific involvement in sensorimotor learning processes. In general, two procedures were used to select appropriate ROIs. First, when investigating a specific localized region, appropriate masks for each ROI were selected based on MNI coordinates in the literature related to sensorimotor processes. Then, to use a more standardized approach that would generalize across studies, a corresponding ROI mask that contained the MNI coordinates and belonged to an established imaging atlas (see Table 2). Five ROIs were from the Harvard-Oxford (HO) Atlas, lateralized and investigated separately (SPL, M1, S1, PMv, LO) and another one from the cerebellar parcellation of the AAL atlas (right and left CBVI). For M1 and S1, the HO atlas ROI was used for the left hemisphere, but for the right, our functionally localized ROIs were used from the GcSS procedure (which incorporated the HO atlas regions). Second, we also wanted to investigate specific networks. Therefore, we used CONN’s identified network regions (see Figure 4). These regions are provided as part of the toolbox and reflect an ICA analysis of the HCP dataset (n=497), and a hierarchical clustering analysis (Bar-Joseph et al., 2001; Sorenson, 1948) applied to the Cambridge 1000-connectomes resting state dataset (n=198; http://www.nitrc.org/projets/fcon_1000). Of the available network ROIs, there were 13 directly relevant to sensorimotor processes, outlined in Table 1. This made five (10 when lateralized) from the HO atlas, one (two when lateralized) from the AAL atlas, and 13 from CONN.

**Figure 4:**
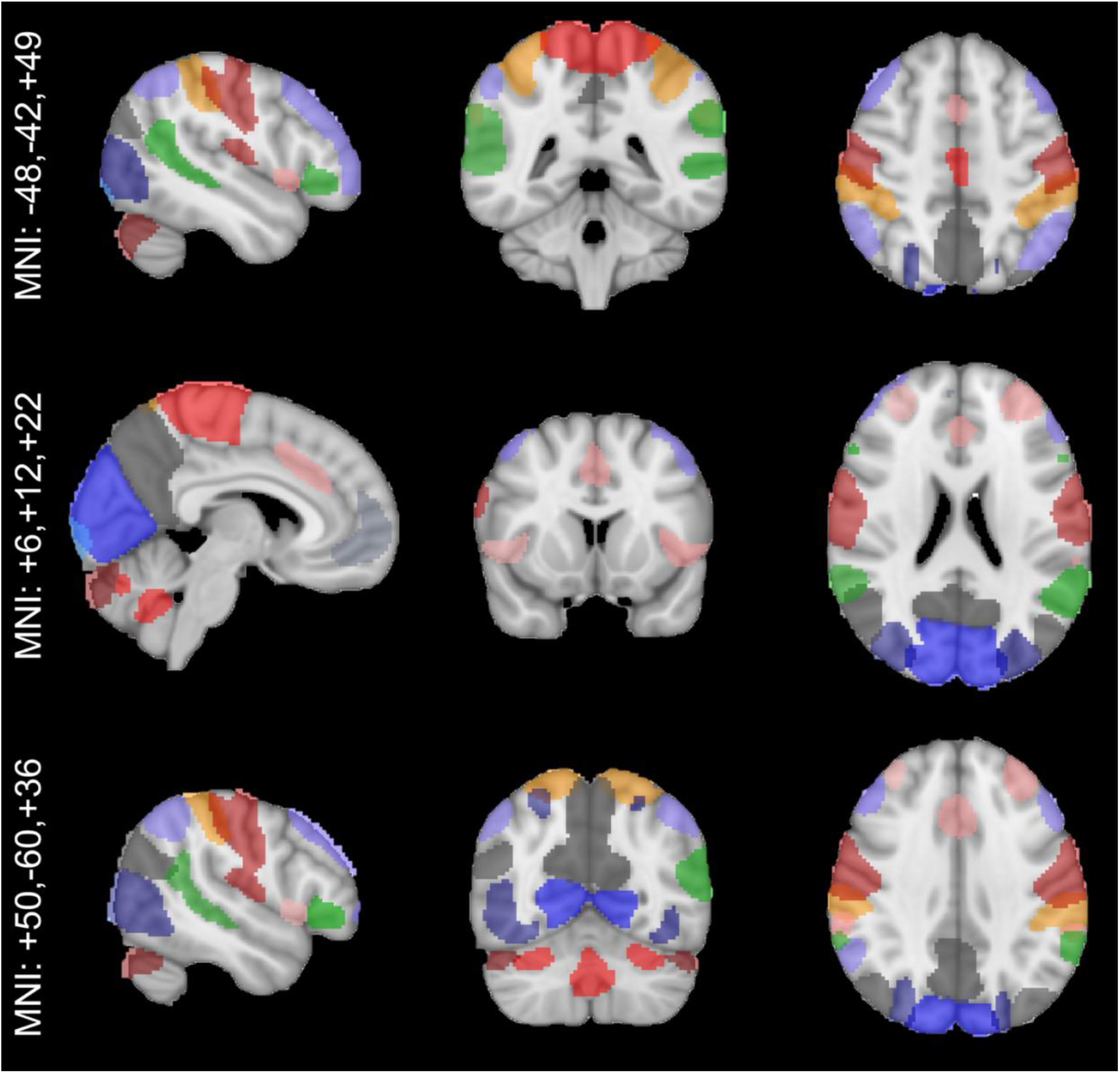
Visualization of the CONN network ROIs provided with the toolbox. Red is SMN and CB network seeds. Blue is VN seeds. Orange is DAN seeds. Purple is CEN seeds. Green is language network seeds. Grey is DMN seeds. Pink is SN seeds.

**Table 2:** Details of the ROIs selected for analysis based on MNI coordinates from the literature, and the primary network(s) each primarily belongs to. Selected ROIs were standard anatomical parcellations from existing atlases that included the target MNI coordinates in a roughly central location.

| ROI | MNI Coordinates | Reference | Atlas | Atlas ROI name | Network |
| --- | --- | --- | --- | --- | --- |
| SMN-LAT | (56,-10,29),<br>(56,-10,29) | (Nieto-Castanon & Whitfield-Gabrieli, 2022) | CONN Analysis | SensoriMotor Lateral | SMN |
| SMN-SUP | (0,-31,67) | (Nieto-Castanon & Whitfield-Gabrieli, 2022) | CONN Analysis | SensoriMotor Superior | SMN |
| VN-MED | (2,-79,12) | (Nieto-Castanon & Whitfield-Gabrieli, 2022) | CONN Analysis | Visual Medial | VN |
| VN-OCC | (0,-93,-4) | (Nieto-Castanon & Whitfield-Gabrieli, 2022) | CONN Analysis | Visual Occipital | VN |
| VN-LAT | (-37,-79,10),<br>(38,-72,13) | (Nieto-Castanon & Whitfield-Gabrieli, 2022) | CONN Analysis | Visual Lateral | VN |
| DAN-FEF | (-27,-9,64),<br>(30,-6,64) | (Nieto-Castanon & Whitfield-Gabrieli, 2022) | CONN Analysis | Dorsal Attention (FEF) | DAN |
| DAN-IPS | (-39,-43,52),<br>(39,-42,54) | (Nieto-Castanon & Whitfield-Gabrieli, 2022) | CONN Analysis | Dorsal Attention (IPS) | DAN |
| CBa | (0,-63,-30) | (Nieto-Castanon & Whitfield-Gabrieli, 2022) | CONN Analysis | Cerebellar Anterior | CN |
| CBp | (0,-79,-32) | (Nieto-Castanon & Whitfield-Gabrieli, 2022) | CONN Analysis | Cerebellar Posterior | CN |
| LO | (-48,-78,6) | (Q. Huang et al., 2022) | Harvard-Oxford atlas (Desikan et al., 2006) | Lateral Occipital Cortex, inferior division | VN |
| SPL | (+10 -52 +60) | (Limanowski & Blankenburg, 2016b) | Harvard-Oxford atlas (Desikan et al., 2006) | Superior Parietal Lobule Right | DAN, SMN |
| S1 | (+50 -21 +52) | (Holmes & Tamè, 2018) | Harvard-Oxford atlas (Desikan et al., 2006) | Postcentral Gyrus Right | SMN |
| M1 | (+38 +15 +58) | (Holmes & Tamè, 2018) | Harvard-Oxford atlas (Desikan et al., 2006) | Precentral Gyrus Right | SMN |
| PMv | (+56 +12 +22) | (Limanowski & Blankenburg, 2016b) | Harvard-Oxford atlas (Desikan et al., 2006) | Inferior Frontal Gyrus, pars opercularis Right | DMN, SMN, VAN |
| CBVI | (-32 -52 -28) | (Vahdat et al., 2011c) | AAL atlas, cerebellar parcellation (Tzourio-Mazoyer et al., 2002) | Cerebelum 6 Left | SN, RN, VAN, SMN, DAN |

The seed-based connectivity was used for subsequent second-level inference in a <u>seed-based analysis</u>. This work explored 25 seeds. First, there were four networks investigated, with seeds defined using the cortical network seeds (Fig. 4) identified from a study of 497 participants (Nieto-Castanon & Whitfield-Gabrieli, 2022). Secondly, regional seeds were chosen from two standard neuroimaging atlases, the Havard Oxford Cortical Atlas (Desikan et al., 2006) and the cerebellar parcellation from the automated anatomical labeling (AAL) atlas (Tzourio-Mazoyer et al., 2002). In addition, two regional seeds were defined by the functional localizer, right M1 and S1. In total there were six regions of interest that were tested bilaterally.

***ROI-to-ROI Connectivity Matrices***. Individual ROI-to-ROI connectivity (RRC) matrices contain the functional connectivity between all pairwise ROI parcels, again computed as the bivariate Fisher-transformed correlation coefficients between all pairwise ROI BOLD timeseries. These matrices were used to inform the second level <u>whole brain network connectivity analysis</u>.

***Graph-Theory Adjacency Matrices***. Binary adjacency matrices were constructed from the ROI-to-ROI connectivity matrices for each subject with an edge threshold of the Fisher-transformed Pearson’s R (z-score) of ±0.15. Connection thresholds for connections from significant ROIs were set as p<.01 (Benjamini & Hochberg, 1995). These adjacency matrices were applied in the second-level <u>graph-theory analysis</u>.

**Second-Level Analyses**. The focus of the current investigation was the whole-brain dynamics related to behavior. To this end, our goal was to both provide context for highly specific and localized regions previously implicated in upper-limb reaches involving sensorimotor processes while also considering the larger functional dynamics of the entire brain. Therefore, we began with identifying specific regions to use as seeds in a whole-brain connectivity analysis that related the functional connections to behavior. The resulting functional connections were then used to identify a behavior-related network, which could then be tested under graph-theoretical approaches.

***General Linear Model.*** This model was applied to the unique connectivity (e.g. SBC, adjacency matrix, RRC) maps discussed in each of the following analyses. This models a the linear association, with ordinary least squares, between our dependent measure, the connectivity values, and our independent measures, (e.g. session-within, and sensory behavior-between) with a General Linear Model, GLM (Nieto-Castanon, 2020).

The resting state data did not involve any task during the MRI data acquisition, even though participants did engage in a behavioral reaching task in between resting state MRI data acquisition sessions. While their performance on the behavioral task was hypothesized to be related to the functional connections and patterns in their neuroimaging data, it was not anticipated that measurement of their sensorimotor behaviors would change their functional activity. Therefore we elected to model the session contrast as an average. Using a group analysis, the GLMs were defined with a between-subjects parameter (computed from behavioral data outside of the scanner), and the within-subjects specification of averaging across both resting state scan sessions. Wilks lambda, defined in the context of a likelihood ratio test that compares our hypothesized model to an unconstrained model (e.g. where connectivity could be any value), is used to evaluate our hypothesis that sensory behavior was related to functional connectivity. Test statistics and p-values were derived from Wilk’s lambda using the default implementation in CONN. In total, four between-subject sensorimotor measures (visual and proprioceptive variance, and position and variance weighting) were tested with a separate GLM for each of the three following analyses.

***Seed-Based Analysis***. Using the first-level SBC maps, the seed-based analyses were performed for each voxel, with first-level connectivity measures at this voxel as dependent variables (one independent sample per subject and one measurement per task or experimental condition, if applicable), and groups or other subject-level identifiers as independent variables. This analysis tests each voxel of the brain in relationship to the given seed region. Voxel-level hypotheses were evaluated using multivariate parametric statistics with random-effects across subjects and sample covariance estimation across multiple measurements. Inferences were performed at the level of individual clusters (groups of contiguous voxels). Cluster-level inferences were based on parametric statistics from Gaussian Random Field theory (Nieto-Castanon, 2020; Worsley et al., 1996). Results were thresholded using a combination of a cluster-forming p < 0.001 voxel-level threshold, and a familywise corrected p-FDR < 0.05 cluster-size threshold (Chumbley et al., 2010). This controlled individual seed-based analyses’ error rates; however because multiple seeds and models were explored, the corrected output from CONN was also corrected in Matlab using the function fdr_bh (Benjamini & Hochberg, 1995). The resulting statistics identified clusters of voxels significantly related to one another, and Fisher-transformed correlation coefficients for the modeled correlation between each seed and cluster were produced. This analysis setting in CONN was referred to as Random Field Theory parametric statistics (Worsley et al., 1996).

Examination of the test-statistic would indicate the direction of the relationship between the functional connections and behavioral measures. Specifically, a positive t-value translated to a positive effect size indicating that the effect of the behavioral measure on the connection was positive. Therefore, a positive t-value indicated that increasing the behavioral measure increased the strength of the functional connection.

Conversely, a negative t-value indicated that increasing the behavioral measure weakened the strength of the functional connection. For seed-based analyses, the t-statistic itself was used to threshold connections using the standard value for cluster-based inferences under Random Field Theory parametric statistics, and therefore, with our sample size of 55, our T threshold was |T(53)|≥3.48, with a cluster size threshold of FDR-corrected-p<0.05, and a voxel threshold of p<.001.

***Graph-Theory***. The resulting seeds and connected regions from the seed-based analysis were then used to constrain the adjacency matrices to make four networks (proprioception, vision, position weighting, and variance weighting). The seed-based analysis was used to identify neural regions related to a particular sensory variable, any region (ROI) not identified for a given sensory measure were not retained in the adjacency matrix for that sensory measure. When a seed region identified a cluster of activation that spanned multiple regions (e.g. M1 and S1), all three regions were included as a node for the built network from an atlas parcellation (not an activity-dependent parcellation). Following this, graph theory was applied to the resulting adjacency matrices to evaluate the network composition. Edges were computed with z-scores using a threshold of 0.15 and a two-sided test. The analysis measures computed were global and local efficiency, closeness centrality, eccentricity, average path length, and degree. Significance was determined with a two-sided distribution using an FDR-corrected alpha of 0.05.

***Network Connectivity***. Lastly, we tested the whole-brain network dynamics related to sensorimotor behavior using the RRC matrices, which reflected the functional connectivity between all regional parcels (all Harvard-Oxford cortical and subcortical regions, the AAL atlas cerebellar parcellations, and our two localized regions) as the Fisher-transformed bivariate correlation coefficient between the timeseries data for each pairwise comparison. These matrices then were tested using parametric multivariate statistics for ROI-based inferences formulated from our GLMs (e.g. the between-subject behavioral effect and the within-subject average of both resting state scans) and corrected at the cluster level using the false-discovery rate (Benjamini & Hochberg, 1995; Strang, 2008). Connection threshold was an uncorrected p-value less than 0.01, and ROIs were thresholded at the FDR-corrected p-value of 0.05 using an MVPA omnibus test (Nieto-Castanon, 2022).

***Summary***. Results are presented individually for each behavioral measure, with the three analyses reported. The <u>seed-based analysis</u> tested the connections between each seed and the whole brain. All resulting connected regions were then used to form atlas-based network nodes to compute <u>graph theoretical</u> network measures. Lastly, we tested the whole-brain <u>network connectivity</u> using ROI-level inferences which identified nodes and edges across the whole brain that were related to the behavioral measure (Benjamini & Hochberg, 1995). Full statistic tables and cluster compositions can be found in the supplemental materials.

## Results

### Seed-Based Analysis

This analysis was performed individually for each of the four sensory measures and tested whether the sensory measure was related to each seed’s functional connectivity to each voxel in the brain. The goal of this analysis was to identify the neural regions related to individual sensory biases. See Table 3 for a summary of seeds and clusters.

**Table 3:** Seed-based analysis results. Some seeds were related to more than one cluster. Each row is one connection between a seed and a cluster. Clusters are identified by region as network membership, and often include many neural substrates. MNI coordinates of the peak location and size in voxels for each cluster are provided. r-Pre and r-Post indicate Pearson’s R for pre and post resting state scan sessions, rounded to two decimals. T indicates the test statistic.

| Contrast | Behavioral Measure | Seed | Cluster | Size (k) | Peak (MNI) | T(53) | r-Post | r-Pre | Cluster Network(s) |
| --- | --- | --- | --- | --- | --- | --- | --- | --- | --- |
| Post + Pre | Position Weighting | CN (P) | 1 | 352 | (-54 +00 +28) | -5.24 | -0.13 | -0.11 | SNM, SN, VAN |
| Post + Pre | Position Weighting | LO (R) | 1 | 306 | (-10 -98 +28) | 4.38 | 0.35 | 0.37 | VN, DAN, DMN |
| Post + Pre | Position Weighting | PMV (R) | 1 | 267 | (+10 -24 +34) | 5.55 | 0.09 | 0.12 | SN, DMN, SMN |
| Post + Pre | Position Weighting | S1 (L) | 1 | 275 | (-22 -94 +20) | 4.01 | 0.03 | 0.05 | VN, DAN |
| Post + Pre | Position Weighting | SMN (L) | 1 | 626 | (-22 -94 +20) | 4.4 | 0.03 | 0.04 | VN, DAN |
| Post + Pre | Position Weighting | VN (L) | 1 | 254 | (-54 -16 +38) | 4.29 | 0.11 | 0.08 | SMN |
| Post + Pre | Proprioceptive Variance | DAN: FEF (L) | 1 | 435 | (-44 +46 +26) | 5.01 | 0.13 | 0.12 | CEN, DMN |
| Post + Pre | Proprioceptive Variance | DAN: FEF (L) | 2 | 346 | (-64 -50 -04) | 5.7 | 0.05 | 0.04 | DMN, VN, SMN |
| Post + Pre | Proprioceptive Variance | DAN: FEF (R) | 1 | 824 | (-02 -60 +52) | 5.49 | -0.02 | -0.02 | DMN |
| Post + Pre | Proprioceptive Variance | DAN: FEF (R) | 2 | 387 | (+20 +28 +36) | 5.34 | 0.00 | 0.01 | CEN, DMN, DAN |
| Post + Pre | Proprioceptive Variance | DAN: FEF (R) | 3 | 301 | (-38 -70 +30) | 5.21 | 0.00 | 0.01 | VN |
| Post + Pre | Proprioceptive Variance | DAN: FEF (R) | 4 | 227 | (+54 -50 +18) | 4.67 | 0.04 | 0.00 | DMN, VAN, VN |
| Post + Pre | Proprioceptive Variance | DAN: IPS (R) | 1 | 393 | (+14 -24 +76) | 4.34 | 0.01 | 0.01 | SMN |
| Post + Pre | Proprioceptive Variance | LO (L) | 1 | 960 | (+08 +08 -18) | 5.3 | -0.02 | -0.06 | RN, DMN, SN, CEN |
| Post + Pre | Proprioceptive Variance | M1 (L) | 1 | 380 | (+50 +40 +08) | 5.31 | 0.12 | 0.07 | CEN, DMN |
| Post + Pre | Proprioceptive Variance | PMV (L) | 1 | 489 | (-28 -14 +70) | 5.47 | 0.11 | 0.11 | SMN, CEN, SN, CEN, DMN |
| Post + Pre | Proprioceptive Variance | PMV (R) | 2 | 339 | (-52 -30 +20) | 4.74 | 0.16 | 0.22 | SMN, VAN, DMN |
| Post + Pre | Proprioceptive Variance | PMV (R) | 3 | 287 | (-34 -10 +20) | 5.18 | 0.09 | 0.09 | SMN, SN, RN, VAN |
| Post + Pre | Proprioceptive Variance | PMV (R) | 4 | 228 | (+24 -34 +56) | 4.76 | 0.01 | 0.04 | SMN, DMN |
| Post + Pre | Variance Weighting | DAN: FEF (L) | 1 | 321 | (-64 -48 -02) | 5.34 | 0.03 | 0.02 | VN, DMN, SMN |
| Post + Pre | Variance Weighting | DAN: FEF (L) | 2 | 304 | (+66 -40 -08) | 4.96 | 0.00 | 0.00 | VN, DMN, SMN |
| Post + Pre | Variance Weighting | DAN: FEF (L) | 3 | 259 | (+56 -48 +44) | 4.78 | -0.06 | -0.09 | DMN, DAN |
| Post + Pre | Variance Weighting | DAN: FEF (L) | 4 | 229 | (+56 +20 +14) | 5.84 | 0.05 | 0.03 | SMN, DMN, CEN |
| Post + Pre | Variance Weighting | DAN: FEF (R) | 1 | 402 | (+00 -58 +52) | 4.83 | -0.01 | -0.02 | DMN |
| Post + Pre | Variance Weighting | DAN: IPS (L) | 1 | 438 | (+56 -46 +40) | 5.28 | 0.02 | -0.02 | DMN, DAN |
| Post + Pre | Variance Weighting | DAN: IPS (L) | 2 | 216 | (-68 -44 +02) | 4.68 | 0.02 | 0.01 | VN, DMN, SMN |
| Post + Pre | Variance Weighting | DAN: IPS (R) | 1 | 301 | (-68 -44 +02) | 4.88 | -0.02 | -0.08 | VN, DMN, SMN, VAN |
| Post + Pre | Variance Weighting | PMV (R) | 1 | 277 | (-60 -38 +12) | 5.33 | 0.13 | 0.18 | SMN, VAN, DMN |
| Post + Pre | Variance Weighting | SPL (L) | 1 | 304 | (-66 -44 +04) | 5.16 | 0.04 | 0.01 | VN, DMN, SMN, VAN |
| Post + Pre | Visual Variance | CN (A) | 1 | 675 | (-28 +10 -04) | -5.79 | 0.14 | 0.13 | SN, RN, VAN |
| Post + Pre | Visual Variance | PMV (R) | 1 | 468 | (-28 -54 -40) | -5.52 | 0.07 | 0.10 | DAN, SMN, VAN, RN, SN, VAN |
| Post + Pre | Visual Variance | PMV (R) | 2 | 290 | (+30 -54 -36) | -5.28 | -0.04 | -0.05 | DAN, SMN, VAN, RN, SN, VAN |
| Post + Pre | Visual Variance | PMV (R) | 3 | 218 | (+36 -90 +10) | 4.28 | -0.09 | -0.04 | VN |

### Visual Variance

Individuals with the lowest visual variance generally had higher functional connectivity between striatal and cerebellar regions and the right PMv, suggesting motor control may reduce visual variance. Two seeds were found to have significant functional connections. First, the anterior cerebellar network seed showed significant functional connectivity to a cluster in the left hemisphere including striatal regions, specifically the left pallidum and putamen (Fig. 5A & Fig. 6A). This negative effect indicated that individuals with lower visual variance were more strongly functionally connected. The functional connectivity was positive, indicating that these regions grew more active or inactive together in association with lower visual variance (greater precision in estimating visual target location). Second, the right PMv showed functional relationships with three clusters.

**Figure 5:**
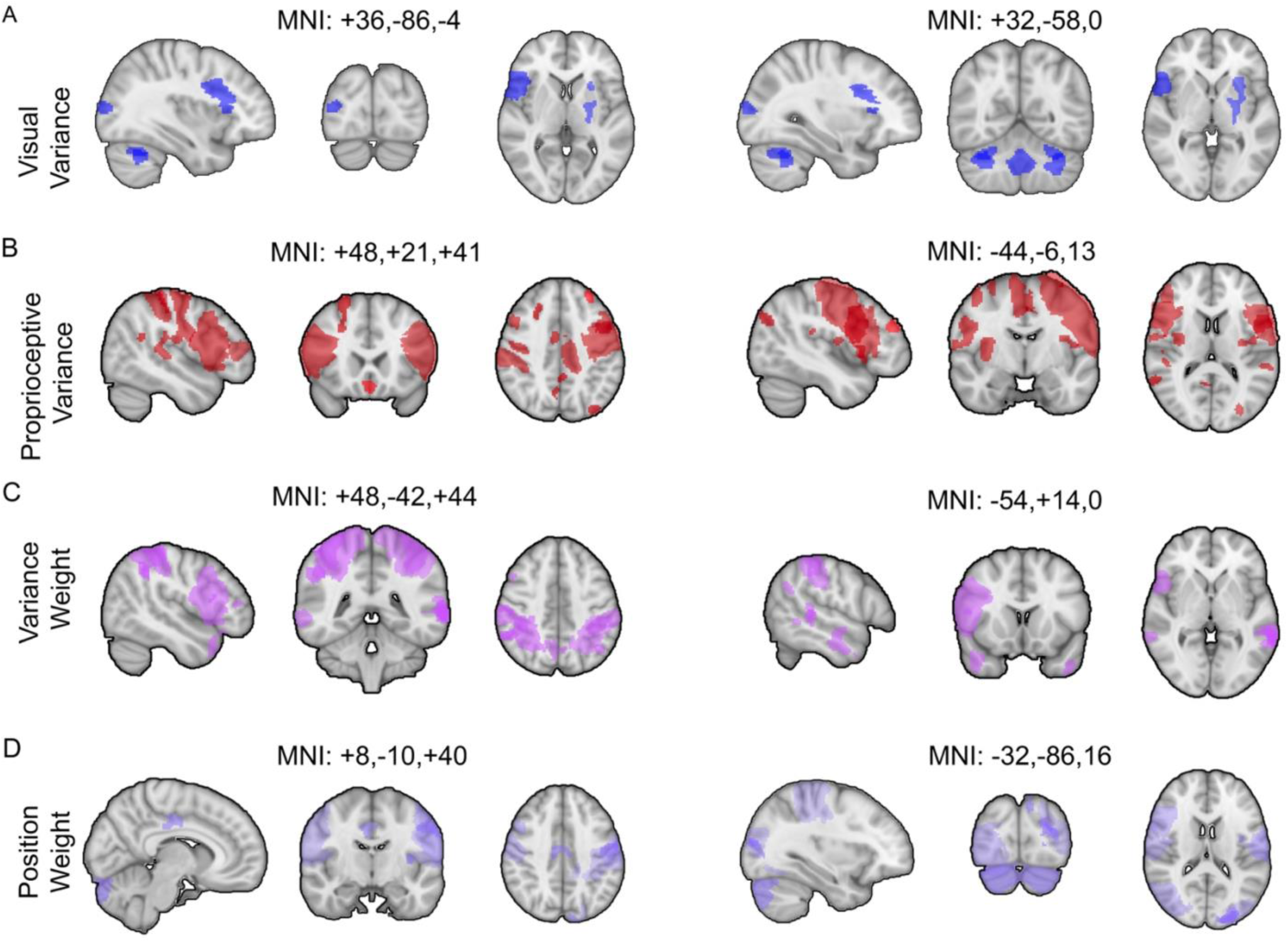
Maps of significantly related seeds and clusters from seed-based analysis. Slice coordinates listed above each set of three slices. Each row corresponds to a sensory measure. Color corresponds to behavioral measure (Blue: Visual, Red: Proprioceptive, Purple-Pink: Variance Weight, Purple-Gray: Position Weight)

**Figure 6:**
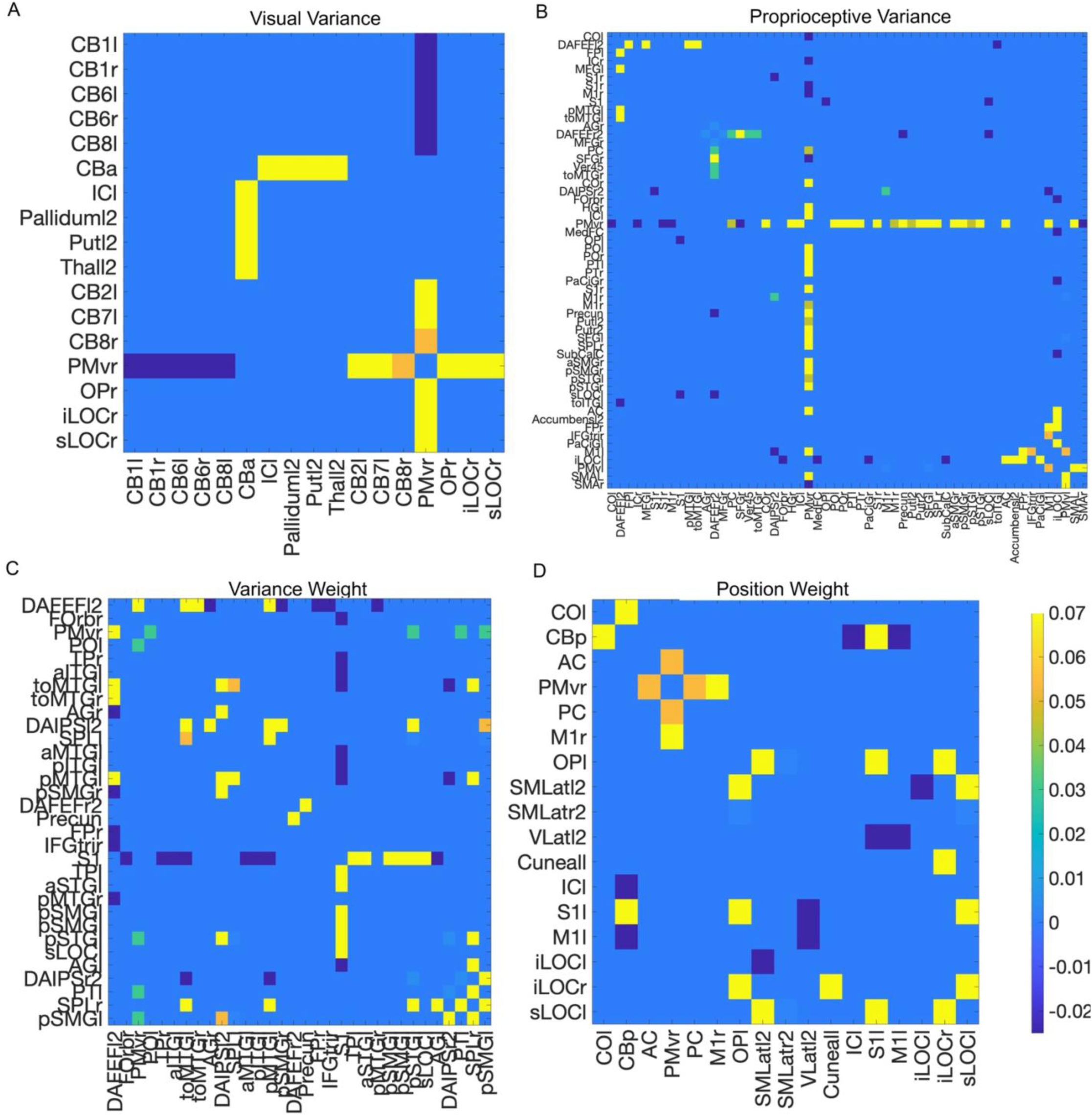
Functional connectivity values from the seed analysis. Panel A: Visual Variance. Panel B: Proprioceptive Variance. Panel C: Variance Weight. Panel D: Position Weight.

Bilateral cerebellar connectivity with the right PMv indicated that lower visual variance was related to increased reciprocal inhibition between these regions. Conversely, lower visual variance was related to increased synchrony between right PMv and left CB II and VII. The right PMv connection to a cluster in the ventral visual stream that included the right lateral occipital cortex indicated that the lowest visual variance was related to the strongest synchrony between these regions.

### Proprioceptive Variance

Individuals with the highest proprioceptive variance were related to widespread functional relationships between sensorimotor, visual, and attentional control regions, including regions of the DMN and regions involved in executive control. Four seed regions had significant functional connections: the DAN seeds, the PMv, M1, and LO. The DAN seeds had widespread connectivity to motor, multisensory, and frontal regions (Fig. 5B & Fig. 6B). The right DAN IPS seed showed connection to a cluster including bilateral M1 and right S1 which indicated that individuals with the highest proprioceptive variance (lowest precision of estimating proprioceptive target location) were related to higher functional synchrony between the DAN IPS seed and the M1/S1 cluster. The right DAN FEF seed was related to four clusters: three multisensory ((1) left dorsal visual stream, (2) right angular gyrus, and (3) the posterior cingulate), and one frontal ((4) superior and middle frontal gyrus). These connections all indicated that higher functional connectivity between these regions was related to higher proprioceptive variance, but the type of connectivity varied. The connection between the DAN and the posterior cingulate indicated that these two regions had higher reciprocal inhibition related to higher proprioceptive variance. The other three cluster connections suggested that when these regions had synchronous activity, proprioceptive variance was highest. Similarly, the left DAN FEF seed was related to two clusters, one a multisensory processing region (middle temporal gyrus), and the other to a region in the central executive network (CEN) (the frontal pole and middle frontal gyrus). These connections suggested that higher proprioceptive variance was related to higher synchrony between DAN FEF and these clusters.

Next, PMv activity had widespread relationships to individual levels of proprioceptive variance. Right PMv showed connections with four clusters largely composed of regions in the SMN and CEN. One cluster included right S1 and SPL, another included the left parietal operculum, and the third included the left central opercular cortex. The fourth cluster spanned a large region including but not limited to right M1, insular cortex, superior frontal gyrus (SFG), supplementary motor cortex (SMA), the putamen, and anterior cingulate cortex. These connections suggested that higher proprioceptive variance was related to higher functional synchrony between these regions. Left PMv also showed significant connectivity to a cluster including left SMA, M1, and SFG indicating that higher proprioceptive variance was related to increased synchrony between left PMv and the cluster. Left M1 also had increased synchrony with the prefrontal cortex, part of the CEN, that suggested lower proprioceptive variance was related to lower functional connectivity between these regions. Lastly, left LO was found to have increased reciprocal inhibition with a medial cluster spanning areas of the RN and DMN that was related to lower proprioceptive variance.

### Variance Weighting

Individuals who weighted vision more than proprioception showed the greatest functional relationships between motor and attentional control regions, as well as with regions responsible for multisensory and sensorimotor integration. In addition, a multisensory hub region was related to the variance weighting network, suggesting the highly integrative and distributed nature of neural activity related to this behavioral measure.

Four seed regions were related to variance weighting, a measure that may be sensitive to many environmental factors like lighting or fatigue. Variance weighting uses variance associated with unimodal estimates to predict how subjects should estimate bimodal targets if they are primarily influenced by variance. Recall that higher values of variance weighting (> 0.5) suggested that multisensory integration was more heavily influenced by vision and lower values of variance weighting (<0.5) suggested multisensory integration was more heavily influenced by proprioception. Thus, a positive effect of variance weighting would indicate higher reliance on vision and a negative effect would suggest higher contribution of proprioception to multisensory integration.

First, the left DAN FEF seed was related to four clusters of activation, and the effect of variance weighting was positive for all four suggesting that higher functional connectivity was related to more reliance on visual information (Fig. 5, row 3). Three clusters were multisensory (bilateral MTG and right angular gyrus (AG) and SMG). One cluster included PMv and prefrontal cortex (PFC). The DAN seed, the bilateral MTG clusters, and the PMv connections suggested that higher synchrony between these regions was related to higher weighting of visual information. Conversely, the connection between the DAN seed and the AG cluster suggested that more reliance on visual information was related to reciprocal inhibition between these regions. The right DAN FEF seed and precuneus were functionally connected related to variance weighting, suggesting that higher reliance on visual information was related to more similar activity between these regions.

Variance weighting also had a positive effect on the left DAN IPS seed, identifying two multisensory clusters, right AG and SMG, and left STG and SMG, whose synchronous activity with the DAN IPS seed was related to higher weighting of vision versus proprioception. The right DAN IPS seed was also positively related to a multisensory cluster including left MTG, which suggested that participants who weighted visual information greater had reciprocal inhibition between these regions. Variance weighting was also related to a functional connection between right PMv and a left-lateralized multisensory cluster including STG, suggesting that higher reliance on visual information was related to increased synchrony between these regions. Lastly, variance weighting was related to a connection between left SPL and a left multisensory cluster including MTG, which indicated that greater reliance on visual information was related increasingly similar activity between these regions.

### Position Weighting

Individuals with higher reliance on vision showed higher functional synchrony between visual, sensorimotor and attentional regions, and reciprocal inhibition related to sensorimotor cerebellar connections. Six seed regions were related to position weighting. Recall that position weight is distributed similarly to variance weight (e.g. high values = more reliance on vision, low values = more reliance on proprioception), but a main conceptual difference is that position weight reflects actual behavior during bimodal trials and may be sensitive to the individual’s attention and goals in addition to the unisensory variances (Block & Bastian, 2011b; Block & Sexton, 2020). First, a functional connection between the posterior CBN seed and a cluster including left M1, S1 and insular cortex suggested that increased reliance on proprioceptive information was related to higher reciprocal inhibition between these regions (Fig. 5, row 4 ). There was also a connection between right LO and a cluster in the VN including early visual cortex that suggested increased reliance on visual information was related to more uniform activity between LO and early visual cortex. There was also a functional connection between right PMv and the cingulate gyrus, which suggested that individuals who gave more weight to visual information (versus proprioceptive) had more similar activity between these regions.

There was a connection between left S1 and a cluster including early visual cortex and regions along the dorsal visual stream, and another connection between the left SMN seed and a cluster including early visual cortex and the dorsal visual stream. Both effects indicated that a higher reliance on visual information was related to more synchronous activity between each seed and cluster. Lastly, the left VN seed was connected to left M1 and S1, which suggested synchronous activity between these regions was related to higher reliance on visual information.

## Graph-Theory

This analysis was tested individually for each of the four sensory measures, and nodes in each adjacency matrix was constrained by the seeds and regions identified in the respective sensory seed-based analysis. The goal of this analysis was to identify network dynamics related to each sensory measure.

### Visual Variance

The visual variance network identified by the seed connectivity analysis included the left striatum (specifically the putamen and pallidum), the insular cortex, bilateral cerebellar lobules I, VI, VIII, the right lateral occipital cortex, and left cerebellar lobules II and VII, almost mirroring regions identified by the seed-based analysis. A test of graph theoretical measures identified no significant nodes or edges (Fig 7A).

**Figure 7:**
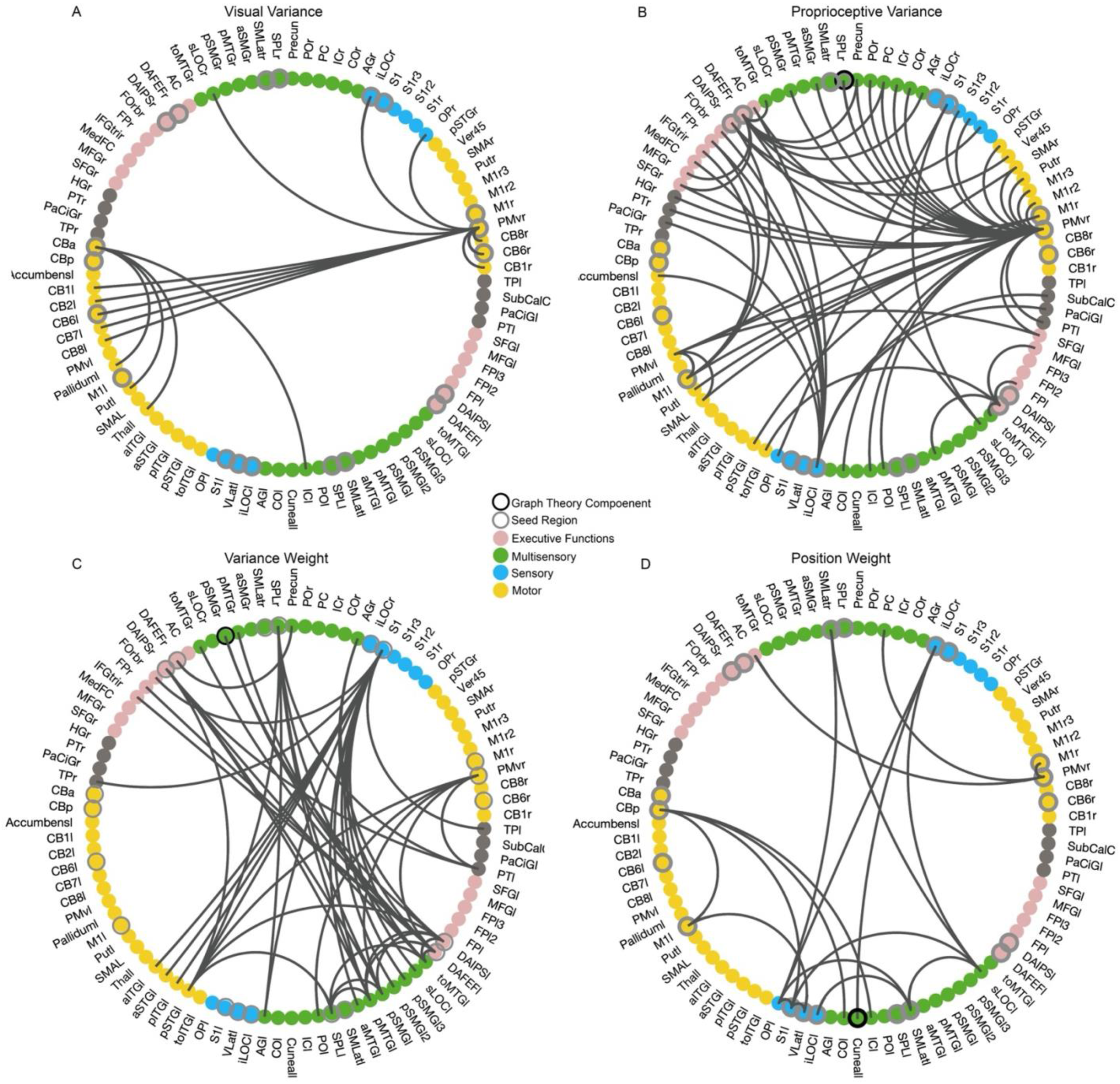
Visualization of the constructed networks in a ring plot sorted by MNI coordinate, right hemisphere to the northwest and left to the southeast. Nodes (circles) in the ring are composed of every region identified across all four sensory measures for ease of visual comparison. Grey circled nodes indicate tested seeds, and black circled nodes signify significant graph-theoretical nodes. Node colors reflect primary function, yellow is motor, blue is sensory, green is multisensory, pink is executive function, and grey is unspecified. Lines signify significant relationships between seeds and clusters (edges). Panel A: Visual Variance. Panel B: Proprioceptive Variance. Panel C: Variance Weight. Panel D: Position Weight.

### Proprioceptive Variance

All identified regions were then applied to construct an adjacency matrix to compute measures of a proprioceptive variance network. In this analysis, the right SPL was identified as a central hub for a proprioceptive variance network. The clustering coefficient was significant which indicated that right SPL tended to be widely interconnected with other nodes in the network (Fig 7B).

### Variance Weighting

All identified regions were then used to inform the nodes in a variance weight network, and the adjacency matrix retained edges that had a Fisher-Transformed R greater than the absolute value of 0.15. This analysis identified the right posterior SMG as a network hub, indicating high global efficiency and closeness centrality for this node (Fig 7C). The global efficiency is a measure of how well information travels throughout the network, and the statistical test suggested that removing SMG would negatively impact global efficiency in this network. Closeness centrality measures how interconnected a node is to other nodes, and the SMG was significantly interconnected. The right SMG node also had low eccentricity and average path length, again suggesting this node as highly interconnected within the network. Eccentricity is a measure of how far a node is from the most distant reachable node, and the negative effect on SMG suggested that this distance was short for SMG. Average path length measures the number of steps between all pairs of nodes, and the negative effect on SMG suggested that there were fewer steps overall to get to SMG from other nodes. Together all these results suggest that SMG is highly interconnected in the variance weight network.

### Position Weighting

These identified regions were then used to construct an adjacency matrix including only these regions as nodes, and then thresholding edges based on Fisher’s Z. A multisensory node, the left cuneal cortex, identified a clique, or small world, for the position weight network (Fig 7D). The cuneal cortex had high local efficiency, suggesting that the information flow of all nodes around the cuneal cortex was very high. The cuneal cortex also had a high clustering coefficient, again indicating that the nodes around the cuneal cortex were inter-connected. Together, these results indicate that connections between multisensory regions were very efficient in the position weight network, and that the cuneal cortex itself was not vital to these connections. This suggested that the goal-related weighting of vision versus proprioception was a dynamic process occurring at the stage of multisensory integration

## Network Connectivity

This analysis was again performed individually for each sensory measure to test its relationship to the functional connections between all ROI-parcels in the whole-brain. The goal of this analysis was to identify the whole-brain network dynamics related to sensory bias, without imposing specific ROI-related hypotheses.

### Visual Variance

The whole brain network analysis of visual variance suggested strong involvement of the dorsal and ventral attention and salience networks (Fig 8A). The right salience network node in the anterior insula (F(2,52)=9.75, p-FDR=.042074) showed functional synchrony with the right hippocampus, bilateral fusiform cortex, and the left DAN FEF node; and reciprocal inhibition with the right posterior superior marginal gyrus and vermis 8.

**Figure 8:**
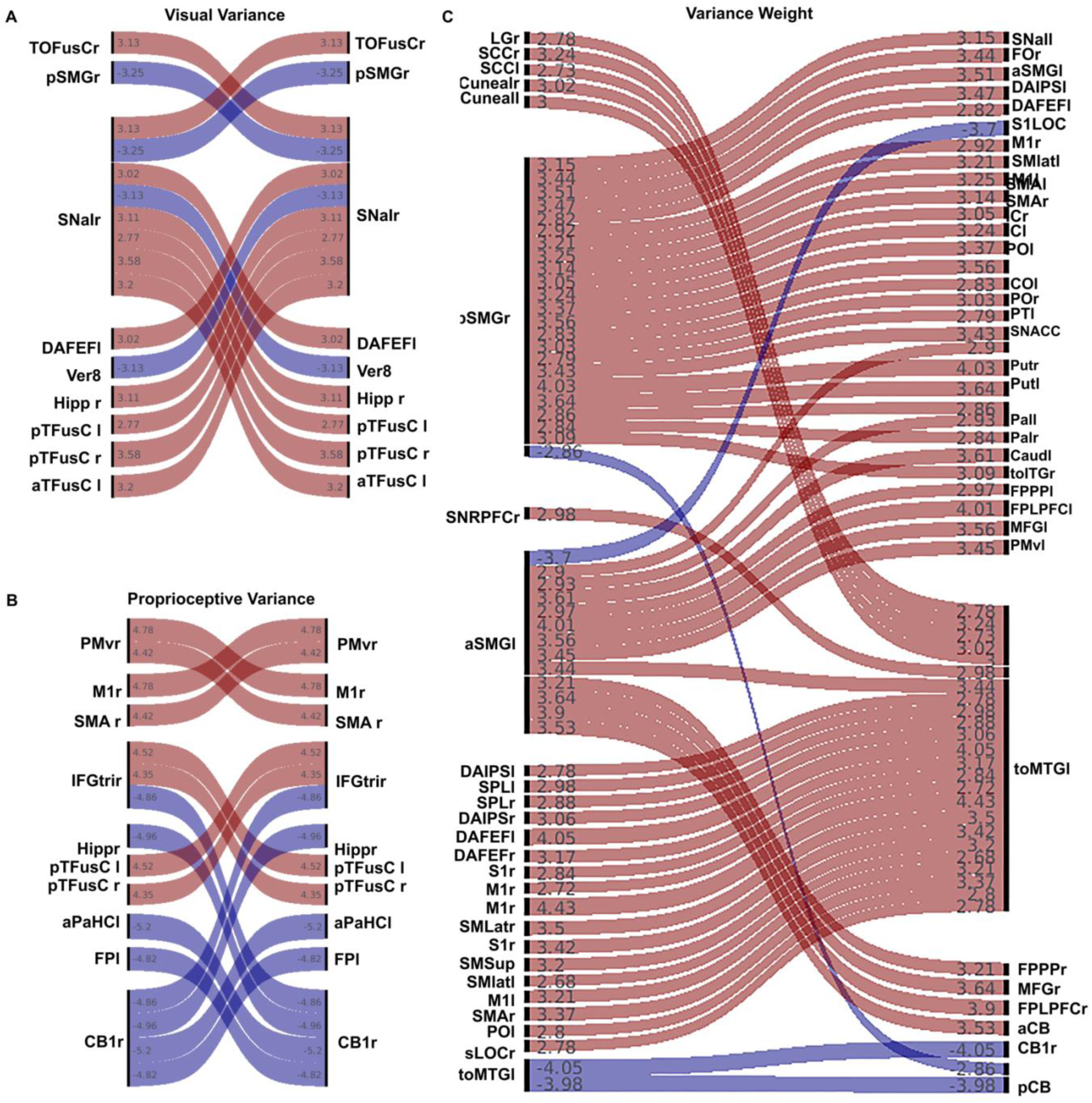
Results from the whole brain network analysis. Panel A: Visual Variance. Panel B: Proprioceptive Variance. Panel C: Variance Weight.

### Proprioceptive Variance

Finally, a whole-brain ROI-level network analysis was computed and 33 ROIs were identified as having connections related to proprioceptive variance (Fig 8B). Due to the over 100 connections resulting from this, we adjusted the connection threshold from p<.01 to p<.0001. This retained the same 33 ROIs but reduced the number of connections to 16. Here, we will discuss the connected ROIs. Activity in right cerebellum crus I (F(2,52) = 17.55, p<.001) showed reciprocal inhibition with the left hippocampus (F(2,52) = 6.70, p=.003), parahippocampal gyrus, and prefrontal cortex related to higher proprioceptive variance. The right PMv had significant positive connections with right M1 and SMA. The right prefrontal cortex was positively related to the bilateral fusiform cortex in the ventral visual stream. Overall, these results support the findings from the seed-based analyses and suggest that proprioception involves widespread recruitment of neural systems including motor, visual, and multisensory systems.

### Variance Weighting

Lastly, a whole-brain ROI-level network analysis was computed which identified three significant ROIs and 64 edges (Fig. 8C). Left MTG (F(2,52)=9.03, p-FDR=0.024) had widespread synchrony with DAN and SMN regions including bilateral SPL, M1, S1, and FEF. Right posterior SMG (F(2,52)=11.39, p-FDR=0.007) had widespread synchrony with DAN, SMN, VAN, and SN regions including left FEF, PMv, IPS, and FEF; and bilateral M1 and striatum; and right S1. Left anterior SMG (F(2,52)=11.27,p-FDR=.007) had widespread positive connections with bilateral striatum, CEN regions, and cerebellum. Anterior SMG also had a negative connection with right S1. Lastly, there were negative connections between cerebellum, SMG, and MTG. This suggested that variance weighting involved attention, sensorimotor, frontal, and salience networks, consistent with the seed-based analysis.

### Position Weighting

Finally, a whole-brain ROI-level network analysis was computed, and there were no significant results.

## Discussion

Here we examined the relationship between visuo-proprioceptive perception during a bimanual pointing task and the whole-brain network dynamics using resting-state functional magnetic resonance imaging. Results suggest that individual sensory biases had systematic influences on functional connectivity. Activity between sensorimotor regions and default mode network (DMN) nodes was related to weighting of vision vs. proprioception. The ventral premotor cortex (PMv) emerged as an important node for sensorimotor integration. Connections from multisensory integration regions like the middle temporal gyrus (MTG) and the superior parietal lobule (SPL), and motor coordination regions in the cerebellum, were related to increased reliance on visual versus proprioceptive information. These findings suggest that individual sensory biases during sensorimotor behavior may be characterized by specific and specialized patterns of neural activity.

### Internal models and control

Individual neural activity patterns have long been recognized in the DMN, a system traditionally associated with self-reflective cognition like future planning, mind-wandering, and consciousness (Bodien et al., 2017; Buckner et al., 2008; Menon, 2023). Current theories of the DMN identify its role in the construction and maintenance of internal models of the self and environment, which directly influence the perception and integration of sensory information (Buckner et al., 2008; Menon, 2023). The adjustment or reweighting of these internal models in accordance with individual sensory perception tendencies (e.g. having more noise in the visual versus proprioceptive system) may be reflected in the activity of DMN nodes such as the posterior cingulate cortex (PCC) and medial prefrontal cortex, which are known for their role in the integration of multisensory and affective cues (Andrews-Hanna et al., 2014; Sridharan et al., 2008). Indeed, DAN, LO, M1, and PMv seeds all showed increased synchrony with DMN regions such as the PCC as proprioceptive variance increased. Similarly, increasing variance weighting (indicating more reliance on vision) increased functional connectivity between DAN and PMv seed regions and the DMN, primarily suggesting increased synchrony, although for some connections this indicated reciprocal inhibition between the DMN and the seed regions (e.g. DAN and precuneus, see supplemental material for full table). In addition, the position weighting network found a DMN regional hub, implicating the importance of DMN activity in weighting and integrating multisensory information. Together, these results suggest that DMN activity may reflect how an individual integrates and weights predicted body states and action outcomes with sensory inputs.

The DMN may provide a connectivity fingerprint for sensory integration with internal models, but accurate sensorimotor control also relies on how this information interacts with the rest of the brain. The PMv seeds (mostly the right) were positively related to all four sensory measures (indicating increased functional synchrony related to increasing values of the behavioral measure), and each were related to connections with SMN. Additionally, PMv connections to DMN were related to variance weighting and proprioceptive variance; connections to the SN, DAN, VAN, and RN were related to visual variance; connections to the VN related to visual variance and variance weighting, and connections to SN related to position weighting and visual and proprioceptive variance. Decreasing visual variance was related to increased functional connectivity between the right PMv, DAN, SMN, VAN, RN, and SN. These widespread relationships mediated by functional connections with PMv highlighted its role for regulating and integrating sensory information between multiple networks to support sensorimotor control. This widespread activation also supports the ideal that sensory region processes not only inform but constrain motor control.

Attentional and cognitive control also have an impact on individual sensory biases as can be seen from the relationship between sensorimotor reaching behavior and the functional connections of the DAN and CEN. The DAN is known to actively prioritize specific sensory inputs necessary for task execution (Corbetta & Shulman, 2002; Vossel et al., 2014), and the CEN is largely known for its role in decision making, working memory, and cognitive control (Seeley et al., 2007). Critically, the DAN plans and coordinates eye and head movements to visually sample the environment and the CEN mediates the flexible weighting of the incoming sensory information based on goals, uncertainty, and/or prior experience (Shadlen & Roskies, 2012). Indeed, our data support this finding that increased synchrony between the DAN and CEN related to increased variance weighting and proprioceptive variance. The functional connections between the DAN and the DMN in this dataset further support this idea, suggesting that the DMN’s internal model is either directing or informing the DAN’s sensory sampling, or vice versa. Interestingly, higher visual variance was related to reciprocal inhibition between the PMv and the DAN, perhaps suggesting a reduced weighting of noisy visual information in these participants. Together, this suggests the role of the DAN and CEN in dynamically adjusting sensory information sampling and integration depending on the internal model, sensory processing, and goals of the individual.

### Prioritization and network communication

Information flow between and within these networks represents an important component in understanding individual sensory bias. Our analyses found a relationship between sensory behavior and known “hub” regions that have been identified in the literature: SPL and the anterior insula (aI). Activity in SPL , known for integration of multisensory information, showed increased synchrony related to higher variance weighting (lower visual noise) with a cluster including regions in DMN, SMN, and VN; and higher proprioceptive variance with PMv (Grefkes & Fink, 2005b). In addition, SPL was identified as a possible hub in our proprioceptive variance network, possibly implicating SPL in the re-weighting of proprioception when forming the multisensory percept. Conversely, the aI is known for regulating whether the DMN, CEN, or SN (sometimes VAN) exert the most influence on the processing of incoming sensory information (Arcurio et al., 2015; Menon & Uddin, 2010; Steimke et al., 2017; L. Uddin, 2014). Individual differences in the functional connectivity and activity of the aI is strongly associated with altered sensory weighting and processing (Arcurio et al., 2015; Hanlon et al., 2014). Our results find insular cortex functionally connected to the anterior and posterior cerebellar network seeds related to visual variance and position weighting, suggesting that individuals with higher visual variance in estimates of hand position had decreased synchrony between anterior CB and the insula, whereas individuals who relied more on vision than proprioception had decreased reciprocal inhibition between posterior CB and the insula. Together, these results suggest that these two hub regions play important roles in regulating information flow and sensory weighting that may influence individual sensory processing, particularly during motor planning and execution.

The anterior cingulate cortex (ACC), a node in the SN, has long been established in the neural network community for its dynamic interactions with the CEN and the DMN while allocating attentional resources to salient stimuli and monitoring conflict, which impact how sensory information is prioritized (Menon & Uddin, 2010; Seeley et al., 2007; Sridharan et al., 2008). The ACC may also play a role in motor planning that uses sensory inputs because our data found a functional relationship between the PMv and ACC related to position weighting and proprioceptive variance, suggesting that the PMv and ACC are desynchronized when proprioceptive information is not reliable. Higher proprioceptive variance was also related to reciprocal inhibition between the ACC and LO, suggesting that the ACC may have been upregulating visual information, perhaps spatial object location (e.g. reach target), to compensate for noisy proprioceptive information. These results implicate the importance of the ACC for directing attentional resources for subsequent action.

Individual sensory bias related to connections between the SN, VAN, and RN. During a reach to a target, the SN prioritizes relevant sensory information, the VAN can rapidly adjust the reach trajectory and attentional resources, and the RN is involved in learning and automation of actions (Corbetta & Shulman, 2002; Grahn et al., 2008; Menon & Uddin, 2010; Ohlendorf et al., 2007; Seeley et al., 2007; Sridharan et al., 2008; Valentin et al., 2007). Our data found that SN synchrony with PMv was related to higher proprioceptive variance and higher reliance on visual information vs proprioceptive (position weighting). RN synchrony with the PMv was also related to higher proprioceptive variance. Conversely, reciprocal inhibition between the PMv and the SN, VAN, and RN was related to higher visual variance. This suggests that the PMv gets fast access to prioritized sensory information. In addition, functional synchrony between the DAN and VAN was related to individuals with more reliable visual information (higher variance weighting), suggesting that the VAN did not need to reorient attention as frequently for these individuals. Also, visual processing regions like LO showed higher synchrony with SN and RN when proprioceptive variance was high, suggesting that visual information was used to guide reaches more for these individuals. Reciprocal inhibition between anterior CB and the SN and RN related to lower visual variance, indicating that the flexible processing that shifted between salience, automaticity, and learning circuits was related to improved visually-guided reaching. Together these results suggested that individuals use adaptive strategies, given their own sensory biases, to optimize action goals characterized by functional connectivity patterns between key motor areas like PMv and CB, and stimulus prioritization systems like the SN, VAN, and RN.

### Subcortical influence

The CB has long been associated with motor coordination, and only recently has evidence emerged suggesting that the CB plays a crucial role in whole-brain communication, providing perhaps the most convincing case that motor learning involves the systematic integration, translation, and regulation of many brain networks (Kawabata et al., 2022). This makes the cerebellum an important component in the weighting of sensory information, perhaps especially when automatic or habitual strategies are unavailable. For instance, functional connectivity between the DAN and CB lobules VII and VIII has been related to cognitive task performance, DAN-CBVI synchrony was related to visuospatial processing and attention, and connections between CB, VN, and RN have been related to the progression of movement disorders (Brissenden et al., 2016; H. Chen et al., 2022; Sako et al., 2021). Our results supplement these findings, particularly considering connectivity differences due to visual variance. Individuals with high visual variance had increased functional synchrony between anterior CB and striatum, and between right PMv and CBI, II, VI, VII, and VIII. Also, higher reliance on proprioceptive information in position weighting was related to increased reciprocal inhibition between posterior CB and SMN and SN. Together, these results support the idea that cortico-cerebellar connections may be fundamentally involved in adjusting reliance on visual cues, perhaps particularly when proprioceptive information is more reliable.

Lastly, the widespread involvement of the visual and sensorimotor networks related to sensory behavior during reaching should not be understated. Both the VN and SMN had regions whose functional connectivity was related to all four of our sensory measures. This widespread recruitment of both visual and sensorimotor networks during reaching emphasizes how visual regions and related networks may also display neural activity related to motor control, especially complex motor control that requires the processing and integration of multisensory cues. For example, catching a baseball requires tracking and predicting the trajectory of a moving object, integrating this visual information rapidly with proprioceptive to flexibly guide body position, while simultaneously updating motor plans. One region that likely mediates this integration is the middle temporal gyrus (MTG), often classified as a hub in the DMN (and sometimes included in the VN), particularly its posterior portion MT+(Davey et al., 2016; Ohlendorf et al., 2007). Therefore, the widespread relationships between individual sensory biases, visual processing, the sensorimotor network, the DMN, and executive control emphasize that sensation and action are inextricably linked.

### Limitations and Future Directions

While this research provides insights into the neural substrates and networks involved in individual sensory bias and sensorimotor control, there are several limitations. A primary constraint is the use of large, anatomically defined ROIs that often encompass multiple functionally distinct subregions. The use of more standardized and generalizable ROIs was an important choice for a preliminary investigation, but subsequent investigation could employ more fine-grained parcellations. This would add specificity to the understanding of the neural dynamics of sensorimotor processing. The insular cortex Harvard-Oxford ROI provides one example of this because while the anterior insula is known for its role in salience detection, the posterior insula plays a role in interoception (L. Uddin, 2014; L. Q. Uddin et al., 2017). Using a more robust parcellation, such as provided by FreeSurfer (Desikan et al., 2006; Destrieux et al., 2010; Fischl, 2004), would more precisely represent the neural networks and substrates involved in sensorimotor behaviors. In addition, there are limitations imposed that are inherent to the analysis of resting state data. The correlational relationships between neural regions do not address information flow and are subject to distortion from factors unrelated to individualized sensorimotor processing network structures, such as head motion and physiological noise (Power et al., 2017).

Also, it should be noted that interpretation of negative functional connectivity should be done with caution. There is growing evidence suggesting that negative functional connections may be related to artifacts that arise during global signal processing, a standard (and necessary) preprocessing step in resting-state functional neuroimaging data (G. Chen et al., 2011). While this may be the case without some external variable to relate to functional connectivity, our analyses explored the relationship between individual sensory perception and functional connectivity and thus may not be subject to the same artifacts. For the sake of transparency and because the primary objective of the current study was to identify all neural regions involved in sensory-guided reaches, we have opted to include effects related to negative functional connections.

Building upon the foundational insights of the present study, future research can address the identified limitations and use experimental manipulations to explore sensorimotor processing. For example, subsequent investigation could use the clusters identified here as seed regions to inform a Granger causality analysis (Shojaie & Fox, 2022), which could establish directionality of neural activity to better understand the information flow for multisensory integration and when internal models exert the most influence on this process. To further characterize individual sensory biases other analytical techniques could be utilized, such as cortical mapping using representational similarity analysis or multivariate pattern analysis (Diedrichsen & Kriegeskorte, 2017; Haxby et al., 2001). Furthermore, while this research established the neural systems associated with individual’s default sensory behavior, future work could manipulate the sensorimotor environment to identify the neural networks involved in sensory recalibration, motor learning, or changes in multisensory integration. A psychophysiological interaction (PPI) paradigm (O’Reilly et al., 2012) could explore the continuous dynamic control of sensory behaviors, and identify the neural activity patterns and substrates involved in online sensorimotor control, such as the updating of internal models or the comparison of internal predictions to sensory prediction errors. Regardless, the current research provides a crucial baseline for these future investigations.

## Conclusion

The results emphasize the relationship between individual sensorimotor control and the complex, dynamic interaction of multiple functionally specialized neural networks. The DMN, SN, VAN, and DAN were related to the reweighting and adjusting sensory information, alongside cortico-cerebellar connections.

Specialized multisensory integration regions such as SPL and MT+ were vital integrators of the prioritized and reweighted sensory inputs. The PMv appeared to be crucially involved in connections with regions containing adjusted sensory information from all involved networks, and then using this to form motor plans or share motor plans that could be used to adjust sensory weighting. These complex and interacting network dynamics provide support for frameworks that suggest sensation and action are not only related but mutually inform, constrain, and adapt to one another. The connectivity fingerprints related to individual sensory behaviors identified here emphasize the importance of considering the systematic dynamics of the whole brain in sensorimotor behaviors.

## Supporting information

Supplemental Results Table

## Author Note

We have no known conflict of interest to disclose. CRediT author roles are reported below. All authors were involved in investigation, methodology, visualization, writing – review and editing. Kess L. Folco had the role of writing – original draft, data curation, formal analysis, formal analysis. Hannah J. Block and Sharlene Newman had the role of conceptualization. Hannah J. Block had the role of supervision.

## Acknowledgments

This research was funded by an R01 grant from the National Institute of Heath, NS112367.

